# How wind and currents shape the drift velocity of macrophytes and macroplastic particles – from experiment to model

**DOI:** 10.64898/2026.03.04.709487

**Authors:** Friederike Gronwald, Zhiyuan Zhao, Rolf Karez, Tjeerd J. Bouma, Florian Weinberger

## Abstract

The post-detachment drifting phase of macrophytes, during which they can be alive, dead, or senescent, plays a crucial ecological and biogeochemical role by influencing long-range dispersal, transporting rafting species, affecting carbon sequestration, promoting blooms, and leading to beaching events. In order to predict the dispersal of macrophytes and macroplastic particles and where they will affect the ecosystem, it is important to be able to model how their drift velocities are influenced by hydrodynamic and aerodynamic factors. In this study, we investigated the drift velocity of macrophytes with diverse morphologies and macroplastic particles in a racetrack flume under different current conditions, in combination with and without wind in the same direction as the water current. Our data show that the drift velocity of macrophytes is highly dependent on their buoyancy and affected by morphological characteristics. Wind increased the velocity of the surface water, which in turn increased the drift velocity of both macrophytes and macroplastic particles. However, wind-induced turbulences reduced the overall effect, especially for macrophytes, which protruded minimally above the water surface in comparison to macroplastic particles. For positively buoyant specimens, an existing particle model was experimentally confirmed to predict macrophyte and macroplastic particle drift velocities reliably, irrespective of shape. For negatively buoyant species, we propose a novel equation to predict drift velocity, incorporating the diverse shapes of macrophytes, as well as their interaction with the bottom. These results represent the first step toward the development of trait-based models that represent macrophytes more realistically in dispersal simulations.

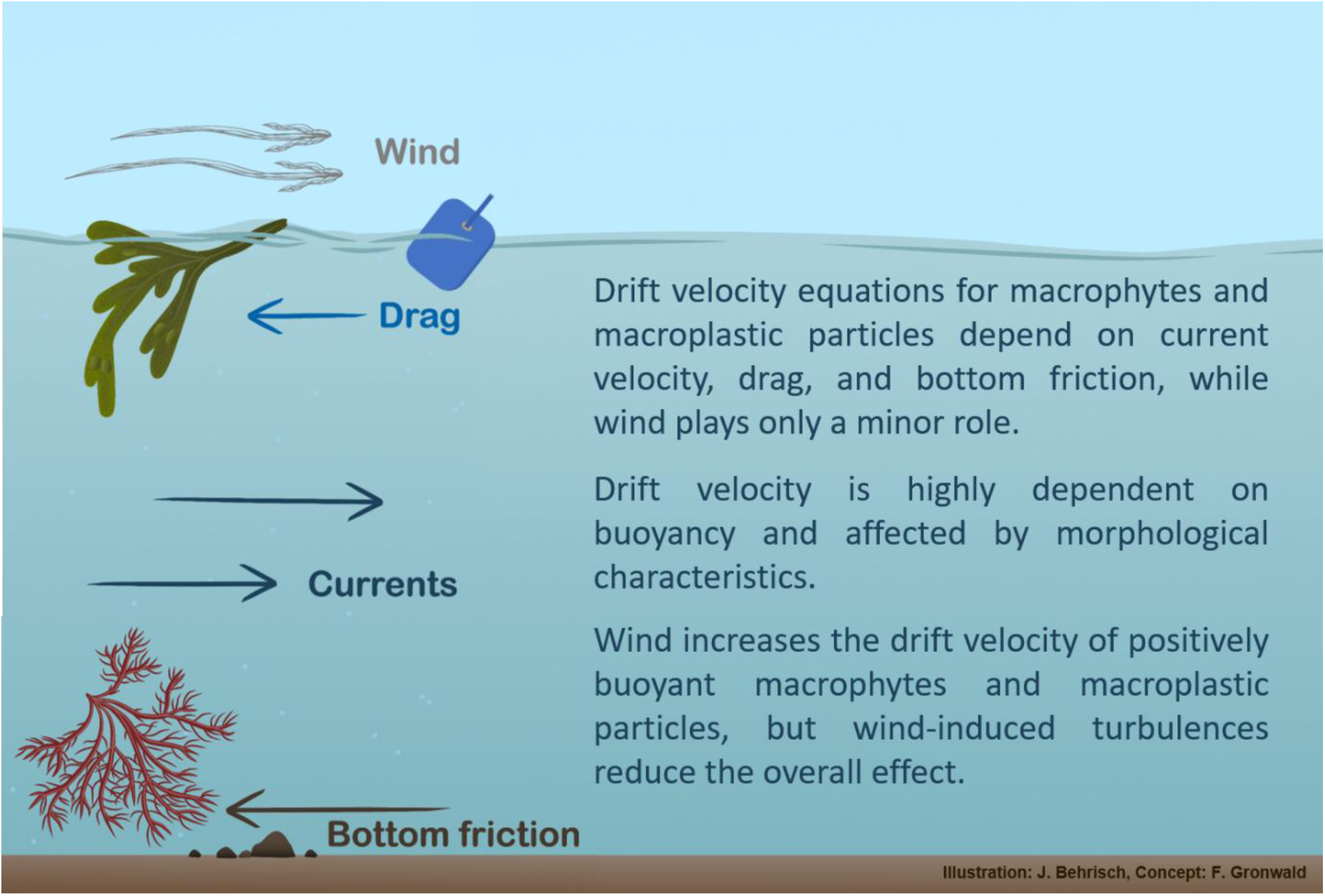

## Introduction

An under-explored aspect of the life cycle of macrophytes is the post-detachment period during which they drift as particles in the ocean. During this period, they may be alive, dead or in various stages of senescence. Regardless, detachment does not end their ecological impact. Much rather, drifting macrophytes contribute significantly to long-range dispersal, facilitating both their own distribution and that of associated rafting species (Holmquist, 1994; van den Hoek, 1987). They can also contribute to carbon sequestration if transported to the deeper ocean. However, drifting macrophytes can also have negative ecological consequences, including the promotion of macroalgal blooms, ultimately leading to increased beaching events (Arroyo & Bonsdorff, 2016; Björk et al., 2023; Ortega et al., 2019). Where these effects occur depends in particular on the drift velocity and trajectory of the macrophytes. These, in turn, are shaped by the hydrodynamic and aerodynamic forces that macrophytes are exposed to while drifting, including currents of variable velocity, tides, waves, and wind (Arroyo & Bonsdorff, 2016; Biber, 2007). Of these, ocean currents and wind in particular determine the speed and direction of movement (Canal-Vergés et al., 2014; Coppin et al., 2024a). Wind directly influences surface water velocity. In particular, within the uppermost layer (0 – 2.5 m) (Bhaganagar et al., 2022; Delpeche-Ellmann et al., 2021) and when aligned with the direction of current flow, wind can enhance surface water velocity.

Among the species that become detached, only a few remain afloat at the water surface due to their positive buoyancy, which can result from density-reducing morphological features, such as gas-filled vesicles in *Fucus vesiculosus* and *Ascophyllum nodosum* or aerenchymatic tissue in *Zostera marina* (Coppin et al., 2024b; Flindt et al., 2004). This buoyancy positions them at the sea-air interface, exposing them to both hydrodynamic drag (influencing the submersed part) and direct wind forcing (influencing the emerged part) (Bhaganagar et al., 2022; Jachowski & Książkiewicz, 2024). In contrast, negatively buoyant species sink and will be mainly affected by hydrodynamic forces.

The focus of research on where and how detached macrophytes drift has so far concentrated on a few ecologically important or nuisance species, such as *Ulva prolifera* in the Yellow Sea (Zhou et al., 2021), *Sargassum* (Bonner et al., 2024), some tropical seagrass species (Lai et al., 2018), and mangrove propagules (Van der Stocken et al., 2015). This has largely overlooked macrophytes with diverging morphologies that actually drift in the ocean and are cast upon our shores (Flindt et al., 2004). In fact, the models developed to predict drift velocities and trajectories have mainly focused on inanimate objects, for example, on life rafts, surface drifters, oil films, icebergs, shipping containers, microplastics, and macroplastics (Christensen et al., 2023; Jachowski & Książkiewicz, 2024; Li et al., 2025; Novelli et al., 2017; Wagner et al., 2022; Wu et al., 2024). A generalized model capable of predicting the drift velocity across a broad range of macrophyte species and other particles is currently missing for broad applications.

In addition to macrophytes, there is also need to understand the drifting behaviour of macroplastic particles. These can have various sizes, shapes, and materials, and can be found drifting in the ocean and accumulating on beaches, posing a major environmental concern (Eriksen et al., 2023). From a physical perspective, macroplastic particles may be regarded analogous to macrophytes, as they share comparable sizes and a similar variety of shapes, and may therefore be influenced in similar ways by hydrodynamic and aerodynamic factors. Consequently, approaches developed to predict drift velocity across a broad range of macrophyte species might be transferable to macroplastic particles as well. Hence, as a secondary objective, we measured the drift velocities of macroplastic particles and evaluated the predictive performance of models developed for macrophytes for macroplastic particles. The easiest method of calculating the drift velocity of a particle in suspension is by assuming that the drift velocity is equal to the current velocity (Flindt et al., 2004; Van der Stocken et al., 2015). For positively buoyant species that are exposed to wind an additional leeway factor of 1 – 4 % of the wind velocity can be added to the sea surface water velocity (Wagner et al., 2022). Since these approaches account for only a limited range of factors, additional models were developed to improve accuracy including factors like inertial effects, as well as lift and added mass forces (Beron-Vera et al., 2019) . Based on the original Maxey-Riley equation (Maxey & Riley, 1983), Beron-Vera et al. (2019) developed a set of equations to predict the drift velocity of finite-sized, buoyant particles drifting on the surface. In Miron et al. (2020) these equations were applied to buoys drifting in a racetrack flume, leading to the verification of the model and a simplification of the equations for predictions in racetrack flume experiments. These sets of equations consider the buoyancy but not the morphological aspects of the particles.

For negatively buoyant macrophyte species, we propose that drift velocity can be calculated based on the water velocity and counteracting effects such as drag and bottom friction. The effect of drag can be expressed as the terminal settling velocity, which can be measured experimentally or calculated theoretically. Gronwald et al. (2025) demonstrated that the sedimentation velocity of macrophytes is influenced by their shape. They proposed a novel approach to determine a shape factor in macrophytes and included it into a drag equation previously developed for ellipsoidal sediment particles (Riazi & Türker, 2019), which allowed them to predict sedimentation velocities of macrophytes with improved accuracy. To our knowledge, bottom friction has not yet been explicitly parameterized for drifting macrophytes, however, formulations for water particles have been proposed by, for example, Dong et al. (2023).

With the aim of developing a general model to predict macrophyte drift velocity, we investigated the drift velocities of positively and negatively buoyant macrophyte species with diverse morphologies in a racetrack flume experiment. We tested the influence of three different water current velocities in combination with and without wind in the same direction as the currents. The results from this experiment enabled us to assess how well existing models predict drift velocities for positively buoyant macrophyte samples and to develop a model for morphologically diverse, negatively buoyant species. Drift velocity measurements were also conducted with macroplastic particles, and the models developed for macrophytes were evaluated for their applicability to macroplastic particles to assess their generality for non-biological particles.

## Materials and Methods

### Sample collection

The macrophytes were collected at several sites in the Baltic Sea (Bülker Weg (54°26’57.4“N 10°11’37.6”E), Schilksee (54°25’16.3“N 10°10’43.1”E), Mönkeberg (54°21’20.92“N 10°10’41.97”E), Schönberger Strand (54°24’40.57“N 10°25’18.45”E), Stein (54°25’7.34“N 10°16’23.20”E) ; ∼ 14 PSU; ∼ 20 °C) around the Kiel Fjord, Germany, as well as at the Osterschelde in Yerseke, the Netherlands (51°30’09.0“N, 4°02’39.7”E), in June 2024. A wide variety of morphologies were collected, comprising a total of 27 different species. The macrophytes collected in the Baltic Sea were transported to the NIOZ in Yerseke (Royal Institute of Sea Research in the Netherlands) within 7 h in cooling bags and otherwise kept in a climate chamber with aeration, light supply and regular water changes (15 PSU, 16 °C) before and during the experiment. Macrophytes collected in the Netherlands were slowly acclimated to 15 PSU by reducing the salinity of the water over several days. See also Gronwald et al. (2025) for details regarding the maintenance of macrophytes. Macroplastic particles of varying shapes and sizes were collected as litter from the beach.

### Drift velocity measurements in the racetrack flume

The drift velocities of the macrophyte species and macroplastic particles were measured in a racetrack flume at the NIOZ. The racetrack flume is a 17 m long and 60 cm wide closed-looped oval filled with 30 cm of seawater (Osterschelde water mixed with tap water, 15 PSU, 17.5 – 19.0 °C). It is equipped with a conveyer belt to generate current velocities, a wave paddle, and an industrial fan that can be added as a wind generator. When using the wind generator, the test section of the flume was completely covered with a transparent sheet to achieve constant wind speeds at the water surface. Water velocity was measured in three dimensions (horizontal, vertical, and lateral) at different water depths using an Acoustic Doppler Velocimeter (ADV, Nortek AS, Norway) at a sampling rate of 200 Hz. The ADV monitors water velocities 5 cm below its sensor head and was coupled to the Vectrino+ programme (-v1.22.00, Nortek AS). After the measurement the values were despiked using the R package (oce), with a window of 50 and a threshold of 1.5. The ADV measurements were verified by direct measurements of the drift velocity of ink pipetted into the water. The surface water velocity was measured using 2.5 x 2.5 cm paper squares that floated on the surface, with five replicates for each treatment. Wind speed was measured with a digital anemometer (Benetech, model: GM8901) at the beginning, middle and end of the test section at three positions each (close to the flume walls and in the middle of the test section) and averaged over the length of the test section.

For the drift velocity measurements, three water current velocities were selected and tested under conditions with and without wind in the same direction as the flow (Table 1). Velocities of positively buoyant samples drifting on the surface and negatively buoyant specimens drifting close to the bottom were measured at all three water velocities. For the wind treatments, the focus was on species drifting on the surface. The drift velocity of each sample was measured five times over the length of a 5 m test section, positioned at the straight part of the flume. The macrophytes were released in the water 1 – 1.5 m before the start of the test section, in order for them to reach terminal drift velocities before starting the measurement. Floating samples were released at the water surface, whereas negatively buoyant samples were released 1 – 2 cm above the bottom. If the samples came in contact with the wall during the measurement the run was cancelled and repeated.

**Table 1.** Water and wind velocities applied in the racetrack flume experiments. Bottom and water column velocity were measured using an ADV, while surface water velocity was determined by measuring drift velocity with paper squares. The numbers in brackets indicate the height (cm) above the bottom at which the water velocity was measured. Bottom water velocities were only measured in the non-wind treatments.

| Treatment | Mean water velocity [m/s] |  |  | Mean wind velocity [m/s] |
| --- | --- | --- | --- | --- |
|  | Bottom (5 cm) | Water column (10 or 15 cm) | Surface (30 cm) | Surface |
| T0.1 | 0.1292 | 0.1462 (10) | 0.1506 | 0.0 |
| T0.1 + wind |  | 0.1448 (10) | 0.1947 | 1.85 |
| T0.2 | 0.2007 | 0.2380 (15) | 0.2344 | 0.0 |
| T0.2 + wind |  | 0.2396 (15) | 0.2691 | 1.85 |
| T0.4 | 0.3990 | 0.4583 (15) | 0.4767 | 0.0 |
| T0.4 + wind |  | 0.4617 (15) | 0.4711 | 1.85 |

[utbl1]

### Morphology and density of macrophytes and macroplastic particles

The morphology of the macrophyte samples was documented by scanning the samples on a flatbed scanner together with a size measure. Resulting images were analysed using Image J (Version 1.54i). The measured parameters included thallus projection area, height and width for macrophytes with flattened morphologies, as well as the width of the 10 smallest branches for macrophytes with filamentous morphologies as described in Gronwald et al. (2025). Positively buoyant specimens were also photographed together with a ruler from different angles through a window pane in the racetrack flume wall while they were floating in non-moving water. To determine how high the negatively buoyant species reach from the bottom into the water column, photos taken through the window pane while drifting were analysed. Each sample was blotted dry and weighed and the volume was measured with the water displacement method. Each measurement was conducted 2 – 3 times depending on the variability of the measurements and the density was calculated with the means. Corey shape factors *Sf* and nominal diameter *dn* were calculated from thallus volumina and thallus size measures as described in Gronwald et al. (2025). For macrophyte species list, densities, shape factors and pictures of all samples see Gronwald et al. (2025). The macroplastic particles were photographed together with a size measure (Supplementary Figure 1).

### Model equations

The drift velocities of specimens were modelled with two different equations that were selected depending on their position in the water column. For species drifting on the surface (positively buoyant) the Maxey-Riley set of equations was used as summarised below in Equation 1 (Beron-Vera, 2020; Miron et al., 2020). For species drifting on the bottom (negatively buoyant) a new equation was developed based on the actual water velocity to which the macrophytes are exposed, the reduction in drift velocity due to drag, and the reduction in drift velocity due to friction with the bottom, as described in Equation 2.

### Drift velocity calculation for positively buoyant species

The drift velocity of floating specimens was predicted according to the Maxey-Riley equation

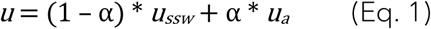

Where *u* is the velocity of the floating particles, *u_ssw_* is the sea surface water velocity, and *u_a_* is the air velocity, which in our experiment was 1.85 m/s for all wind treatments and 0 m/s for treatments without wind. *α* is a buoyancy-dependent leeway factor (Miron et al., 2020) and was calculated as proposed by Beron-Vera (2024). *α* is based on the particle buoyancy *δ*, which depends on particle density 𝜌_𝑝_in relation to the density of the water 𝜌_𝑠𝑤_, with *δ* ≥ 1. It is further assumed that the air-to-water viscosity ratio is *γ* ≈ 0.0167, as suggested by Miron et al. (2020). Water mass density 𝜌_𝑠𝑤_ and kinematic viscosity 𝜈 were calculated after Sharqawy et al. (2010), based on multiple daily measurements of water salinity and temperature in the racetrack flume.

The prediction of *u* for macrophytes in treatments with wind did not fully capture the observed drift velocities (see Results). For this reason, different approaches were designed to disentangle wind-induced changes in surface water velocity from direct wind forcing. The following approaches were tested using Equation 1:

Method 1 (u1): As the equation is intended. For *ussw* the respective surface water velocity was used and for *ua* the respective wind velocity of 1.85 m/s.

Method 2 (u2): For *ussw* surface water velocity with wind was used, but *ua* was 0 regardless of whether the wind generator was used in the treatment or not.

Method 3 (u3): For *ussw* the surface water velocity without wind was used (for example for T0.1 + wind the surface water velocity of T0.1 was used, see Table 1) and for *ua* 1.85 m/s was used. A commonly used and simplified approach to predict drift velocities for specimens floating on the surface involves calculating drift velocity as the sum of sea surface water velocity and a certain percentage of wind velocity, usually in the magnitude of 1 – 4 % (Putman et al., 2018; Wagner et al., 2022). Because macrophytes fall on the smaller end of the drifting particles evaluated so far, we tested this method using an additional 1% of the wind velocity. We refer to this as the simplified leeway-factor method.

### Drift velocity calculation for negatively buoyant species

To predict the drift velocity 𝑢 for negatively buoyant species Equation 2 was developed:

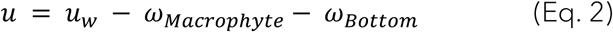

Where *u_w_* denotes the water velocity, 𝜔_𝑀𝑎𝑐𝑟𝑜𝑝ℎ𝑦𝑡𝑒_ accounts for the water drag exerted on the macrophytes, and 𝜔_𝐵𝑜𝑡𝑡𝑜𝑚_ accounts for the interaction between the macrophytes and the substrate. As indicated in Table 1, *u_w_* varies with water depth and decreases notably close to the bottom. We therefore determined *u_w_* at the highest point the macrophyte reaches into the water column (ℎ_𝑝_), with ℎ_𝑝_ measured from the bottom. To be able to calculate *u_w_* at depths between the measured water velocities, an empirical equation was developed. The surface water velocity (𝑢_𝑠𝑠𝑤_) was chosen as reference water velocity, as it provides a consistent basis for scaling velocity profiles. The measured water velocities showed a nonlinear relationship (see Supplementary Figure 2, blue dots), so to determine the best-fitting function, various higher-order polynomial and logarithmic models were tested. Model selection was based on the Akaike information criterion (AIC), resulting in Equation 3, which describes the relationship between horizontal water velocity and depth most accurately in the racetrack flume:

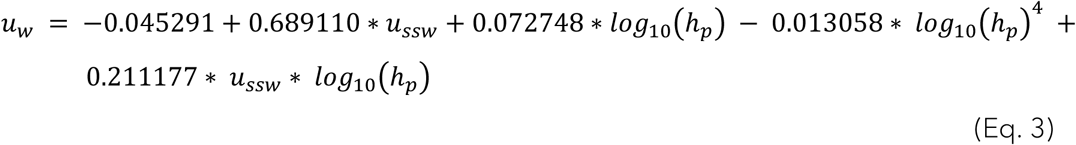

In Equation 2, the terminal sedimentation velocity, which can be determined experimentally or calculated based on the equation developed by Gronwald et al. (2025), serves as an empirical proxy for drag 𝜔_𝑀𝑎𝑐𝑟𝑜𝑝ℎ𝑦𝑡𝑒_.

𝜔_𝐵𝑜𝑡𝑡𝑜𝑚_ accounts for the loss in drift velocity caused by friction between the macrophytes and the substrate and was derived through a multiple step process.

First, to quantify the loss of drift velocity due to friction with the bottom, the experimentally observed drift velocity (𝑢) was subtracted from the preliminary drift velocity calculated without bottom friction (preliminary drift velocity: 𝑢_𝑝_ = *u_w_* − 𝜔_𝑀𝑎𝑐𝑟𝑜𝑝ℎ𝑦𝑡𝑒_). This was done to identify the fraction of drift velocity reduction that could not be explained by macrophyte drag, which we attributed to be the bottom friction (𝐵𝑜𝑡𝑡𝑜𝑚 𝑓𝑟𝑖𝑐𝑡𝑖𝑜𝑛 = 𝑢_𝑝_ − 𝑢 ; Figure 1A pink points).

**Figure 1.**
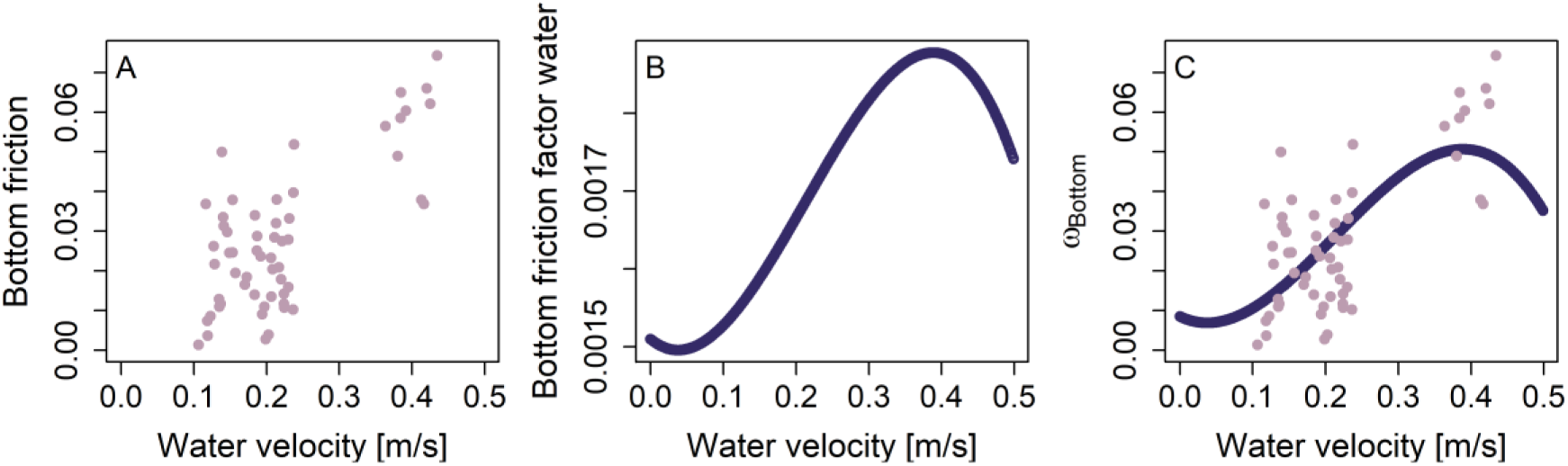
Development of the bottom friction for macrophytes (𝝎_𝑩𝒐𝒕𝒕𝒐𝒎_). A. Calculated bottom friction for each macrophyte sample, pink points. B. Bottom friction coefficient *BFC* after Dong et al., 2023 for water velocities < 0.5 m/s (Eq. 4). C. 𝜔_𝐵𝑜𝑡𝑡𝑜𝑚_ as derived from BFC and used in Eq. 2, together with observed bottom friction values.

The closest bottom friction factor that could be found in the literature is that of Dong et al. (2023), who developed a nonlinear empirical formula for the bottom friction coefficient (*BFC*) for water, which is dependent on current speed (Eq. 4, Figure 1B). For water velocities below 0.5 m/s Dong et al. (2023) defined the current-speed-dependent *BFC* for water as:

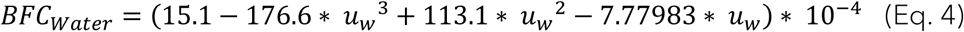

The *BFC* values for water follow a comparable shape, but differ in magnitude from the bottom friction values for macrophytes. This can be seen in Figure 1 when comparing the two scales of the bottom friction (y-axis) of panel A for macrophytes and panel B for water. The average bottom friction for macrophytes is 12 – 31 times higher than that for water, depending on the water velocity.

Finally, to be able to calculate the *BFC_Algae_* for macrophytes without the necessity of experimental data the *BFC_Water_* was scaled to the observed total friction of the macrophytes (Figure 1C). This could be achieved by using linear regression (**𝐵𝐹𝐶_𝐴𝑙𝑔𝑎𝑒_ = 𝑎 ∗ 𝐵𝐹𝐶_𝑊𝑎𝑡𝑒𝑟_ + 𝑏**), with the values for bottom friction for each macrophyte serving as reference values (Figure 1A red points). Consequently, two scaling parameters were obtained a = 114.1662 and b = – 0.1638952.

### Statistics

All plotting and statistical analyses were performed with Rstudio (Version 2023.06.1+524). A Shapiro-Wilk test was performed first to test for normality, followed by a Fligner-Kileen test to test for homogeneity of variance. If the data were not normally distributed and had heterogeneous variances, a Kruskal-Wallis test was performed and, if significant, a pairwise Wilcoxon rank-sum post-hoc test was conducted. This approach was used to test: 1. Whether the drift velocity of the samples differed significantly between treatments, both combined and for each sample individually, 2. whether the drift velocity of the samples differed significantly within each treatment, 3. whether the position in the water column had a significant effect, 4. whether there were significant differences within each position for each treatment, 5. whether wind had a significant effect on the drift velocity. To calculate the first and third quartile the summary function in R was used.

To test whether the drift velocity of the samples differed from the surface water velocity for all treatments with and without wind, the drift velocity was subtracted from the surface water velocity and the result compared to 0. If the differences were normally and homogeneously distributed (Shapiro-Wilk test and Fligner-Kileen test, respectively), a One-sample t-test was performed, if the data were not normally distributed a Wilcoxon signed-rank test was used. A Spearman rank correlation coefficient test was performed to test how the buoyancy-dependent leeway factor α and the buoyancy δ correlate with each other.

To test whether density was significantly different between samples and positively and negatively buoyant species a Shapiro-Wilk test was used to test for normality followed by an ANOVA (analysis of variance).

The model performance for negatively buoyant species, as described by Equation 2, and positively buoyant species, using different Maxey-Riley set of equations and the simplified leeway factor method, was assessed by comparing predicted and observed mean values using median squared deviation (MSD) and Pearson’s correlation coefficient (*r*). To assess linear correlations, Pearson’s correlation coefficient was used; however, because the data was not normally distributed, tested by a Shapiro-Wilk test and histograms, Spearman’s rank correlation coefficient (*ρ*) was used to validate the significance of the models.

All box plots are to be read as follows: the median is represented by the middle line, the box shows the interquartile range between the 25^th^ and 75^th^ percentiles, and the whiskers extending from the box indicate the range within 1.5 times the interquartile range, all points plotted individually represent outliers.

## Results

### Influence of water velocity on macrophyte and macroplastic particle drift velocity

**Figure 2.**
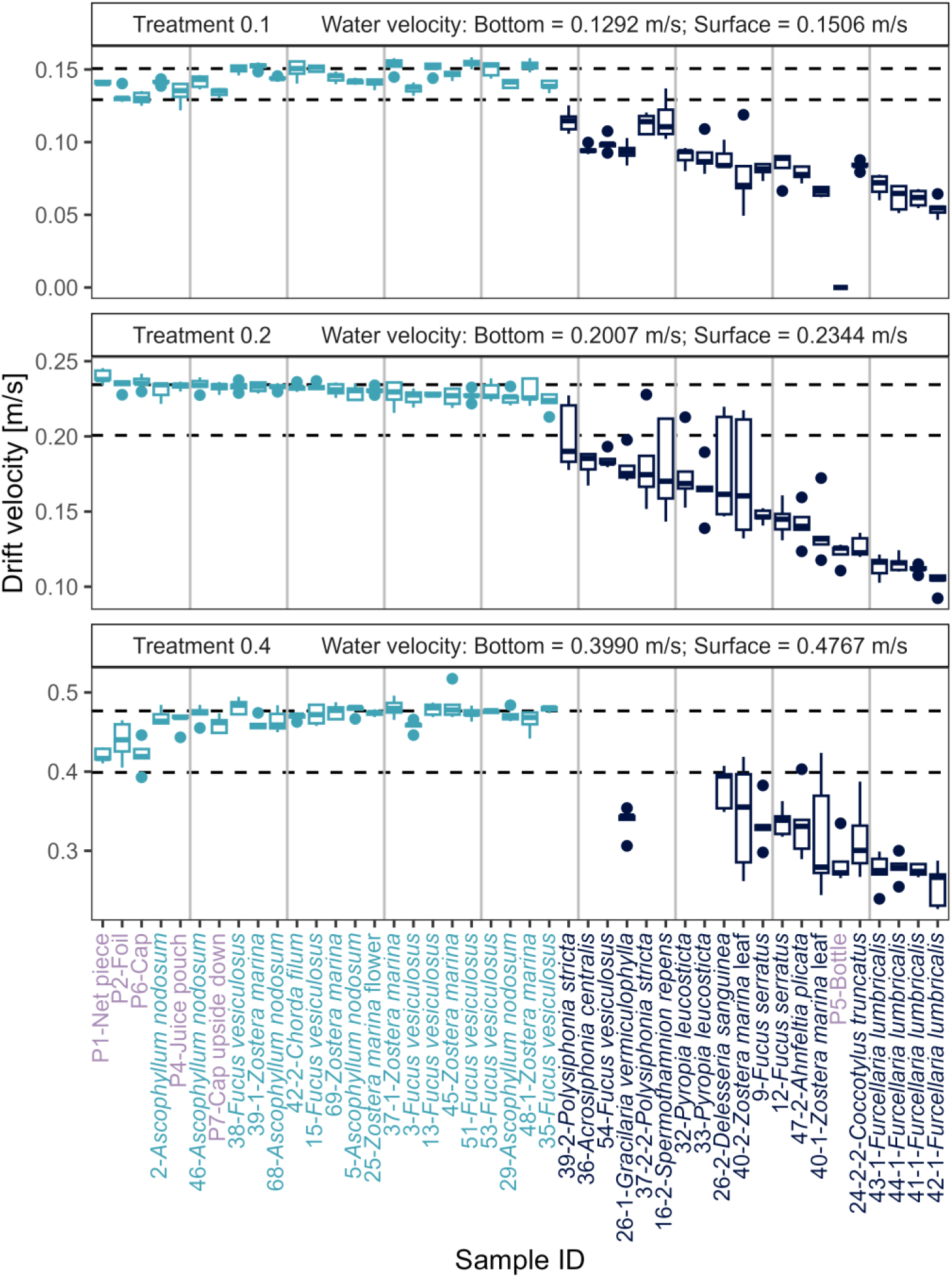
Drift velocity of macrophyte specimens and macroplastic particles. Shown at three different water velocities, coloured according to their position in the water column, positively buoyant in light blue, negatively buoyant in dark blue. Macroplastic particles, which start with ‘P-‘, are indicated by lilac text. Sorted by median drift velocity of treatment 2. In each facet, the upper dashed lines represent the surface water velocity, the lower dashed line represents the bottom water velocity. Only specimens measured in at least two treatments are shown. Vertical lines for orientation.

Overall, a significant difference was observed between the samples at all three water velocities (X² = 600.08, df = 2, p< 0.001). The drift velocity of the samples increased linearly with increasing water velocity (Supplementary Figure 3) and was significantly different for each sample when compared between water velocities (p < 0.05). For all three water velocity treatments, the species differed significantly in their drift velocities (T0.1: X² = 229.53, df = 47, p < 0.001, T0.2: X² = 306.63, df = 68, p < 0.001, and T0.4: X² = 175.9, df = 39, p < 0.001). Observably, all *Furcellaria lumbricalis* (n = 4) samples clustered together at the slower end for all three water velocities, unlike the other species which did not have similar drift velocities within each species. The samples could be divided into two distinct groups according to their position in the water column, either drifting on the bottom or floating at the surface. Species at the surface generally drifted faster than those on the bottom. The position of each sample remained consistent across treatments, i.e. no resuspension occurred for samples drifting on the bottom, even at increasing water velocities. Significant differences in drift velocity between the positions bottom and surface could be observed for each velocity treatment (T0.1: X² = 180.82, df = 2, p < 0.001, T0.2: X² = 247.64, df = 2, p < 0.001, T0.4: X² = 142.86, df = 2, p < 0.001). The drift velocity of samples floating on the surface was more consistent, whereas the drift velocity of samples drifting on the bottom was more variable between samples (Figure 2). Nonetheless, there were significant differences among samples within each position for each velocity treatment (Supplementary Table 1). Also, species drifting on the bottom exhibited a higher within-sample variability and larger standard deviations than floating samples. The drift velocity of the macroplastic particles was of similar magnitude to that of the positively buoyant macrophytes.

A significant effect of density is visible, as density separates the samples into positively buoyant species floating on the sea surface and negatively buoyant species drifting on the bottom (Df = 1, F-value = 300.9, p-value < 0.001, Supplementary Figure 4). However, within the two groups drift velocities are not homogenous and cannot be explained by differences in density. In general, the density between the samples is significantly different when comparing all samples (Df = 63, F-value = 3.299 * 10^28^, p-value < 0.001), and when comparing samples within the negatively buoyant (Df = 36, F-value = 6.829 * 10^28^, p-value < 0.001) and positively buoyant groups (Df = 26, F-value = 1.816 * 10^29^, p-value < 0.001).

In the absence of wind, the drift velocity of floating samples closely matched the surface water velocity across all three treatments. In contrast, bottom-drifting species exhibited slower drift velocities than the water velocity in the bottom layer for the majority of samples. The greatest mean velocity reduction occurred at the lowest water velocity (T0.1), with a 57.79 % decrease, followed by T0.2 (48.52 %), and T0.4 (35.77 %). Interestingly, some species drifted faster than the maximum bottom water velocity, exceeding it by 17.4 % in T0.2, 5.09 % in T0.4, and showing no increase in T0.1. No considerably different speed reductions in red, green, brown macroalgae or seagrasses could be detected, although the highest reductions occurred for the red macroalgae *F. lumbricalis* and *Coccotylus truncatus*.

### How the addition of wind alters the drift velocity of macrophytes and macroplastic particles

A wind velocity of 1.85 m/s in the same direction as the water velocity led to an increase in surface water velocity (Table 1). The increase in surface water velocity decreased with increasing water velocities, with differences between T0.1 + wind and T0.1 = 0.0441 m/s, between T0.2 + wind and T0.2 = 0.0347 m/s, and between T0.4 + wind and T0.4 = - 0.0056 m/s (Table 1). The negative difference between T0.4 + wind and T0.4 is probably due to instrument variability. Within the water column at 10 and 15 cm above the bottom corresponding increases of water velocity due to wind impact were not observed. However, wind also increased the range of possible vertical water velocities at 20 cm above the bottom for the lowest horizontal water velocity (T0.1) by a factor of approximately 3.5, as indicated by an increase in the interquartile range (Table 2, Supplementary Figure 5). However, the vertical water velocity remained constant at higher water velocities, regardless of wind.

**Table 2.** Variability in vertical water velocities. Measured with an ADV at 20 cm above the bottom for each treatment with and without wind.

| Treatment | First quartile (Q1) | Third quartile (Q3) | Interquartile range<br>(IQR = Q3 - Q1) |
| --- | --- | --- | --- |
| T0.1 | - 0.004 | 0.014 | 0.018 |
| T0.1 + wind | - 0.03 | 0.038 | 0.068 |
| T0.2 | - 0.003 | 0.021 | 0.024 |
| T0.2 + wind | - 0.006 | 0.015 | 0.021 |
| T0.4 | - 0.013 | 0.01 | 0.023 |
| T0.4 + wind | - 0.016 | 0.006 | 0.022 |

The drift velocity of the macrophytes species and macroplastic particles changed significantly when wind was applied (for T0.1 vs. T0.1 + wind: X² = 149.33, df = 1, p < 0.001, for T0.2 vs. T0.2 + wind: X² = 115.81, df = 1, p < 0.001, and for T0.4 vs. T0.4 + wind: X² = 23.87, df = 1, p < 0.001). This was particularly evident for the slower water velocities T0.1 (+ wind) and T0.2 (+ wind), whereas for T0.4 (+ wind) the differences were smaller. The addition of wind resulted in an increase in drift velocity for treatments T0.1 + wind and T0.2 + wind for all samples, while drift velocities slightly decreased in T0.4 + wind for macrophytes. However, macroplastic particles showed an increase in drift velocity across all three treatments, including T0.4 (Figure 3).

**Figure 3.**
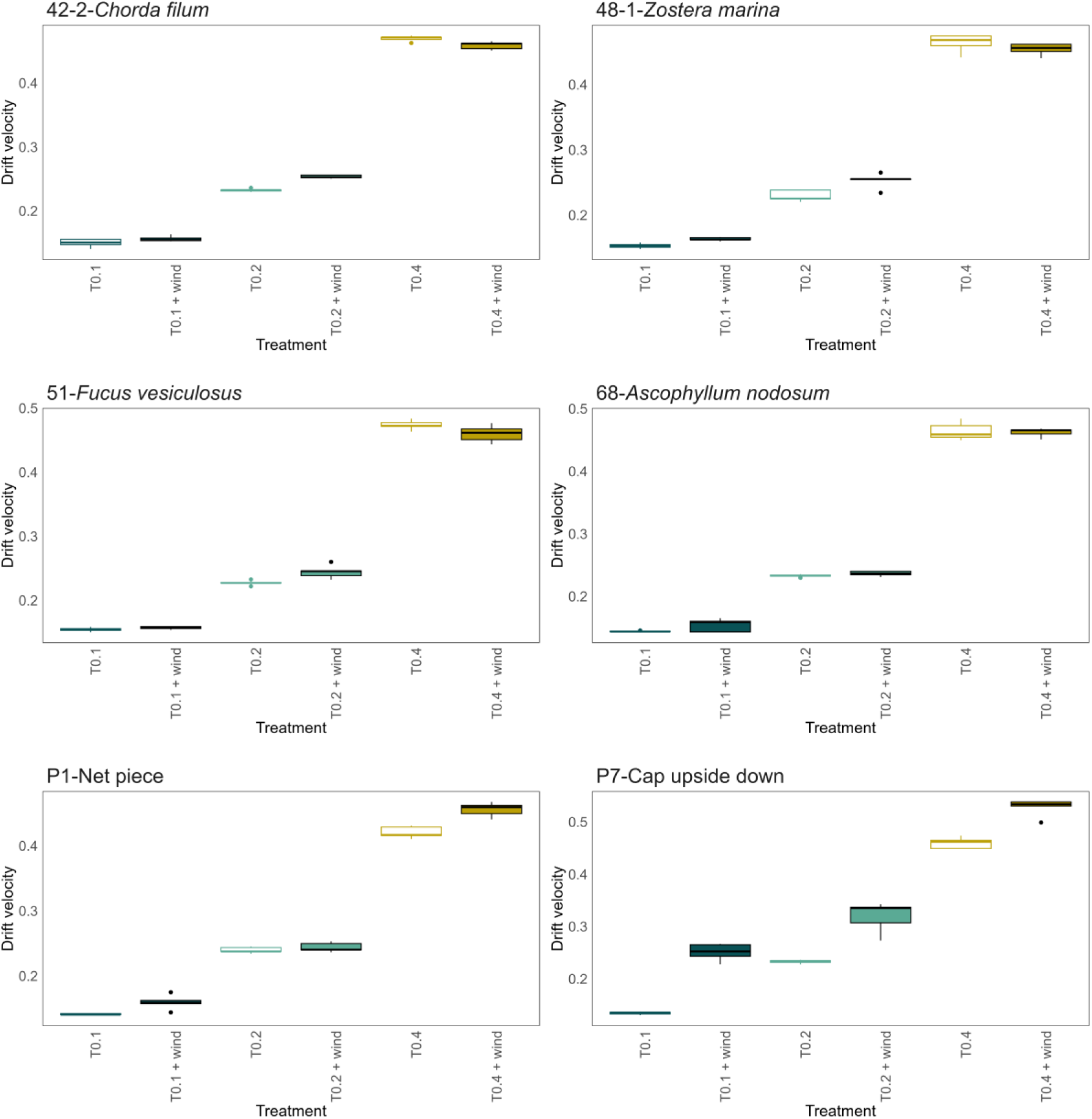
Drift velocities for selected positively buoyant macrophytes and macroplastic particles. Individually depicted for the three current velocities with and without wind. The figure showing all 24 comparisons can be found in the supplements (Supplementary Figure 8). P7-Cap upside down denotes a bottle cap floating with its closed side flat on the water surface.

Thus, while wind resulted in an increased surface water velocity a corresponding increase of drift velocities of floating macrophytes was not always observed. This gets apparent when we look at the differences between surface water velocity and macrophyte drift velocity. The drift velocity of the macrophytes in the treatments without wind is closer to the surface water velocity (Figure 4), indicated by a value close to 0, in comparison to treatments with wind, even though the drift velocities in all treatments differ significantly from the surface water velocity (Supplementary Table 2). The lower the water velocity to which the wind velocity was added the higher the difference between the surface water velocity and the drift velocity, indicating that a slower water velocity at the same wind speed allows for less increase in drift velocity of the macrophytes in relation to the water velocity.

**Figure 4.**
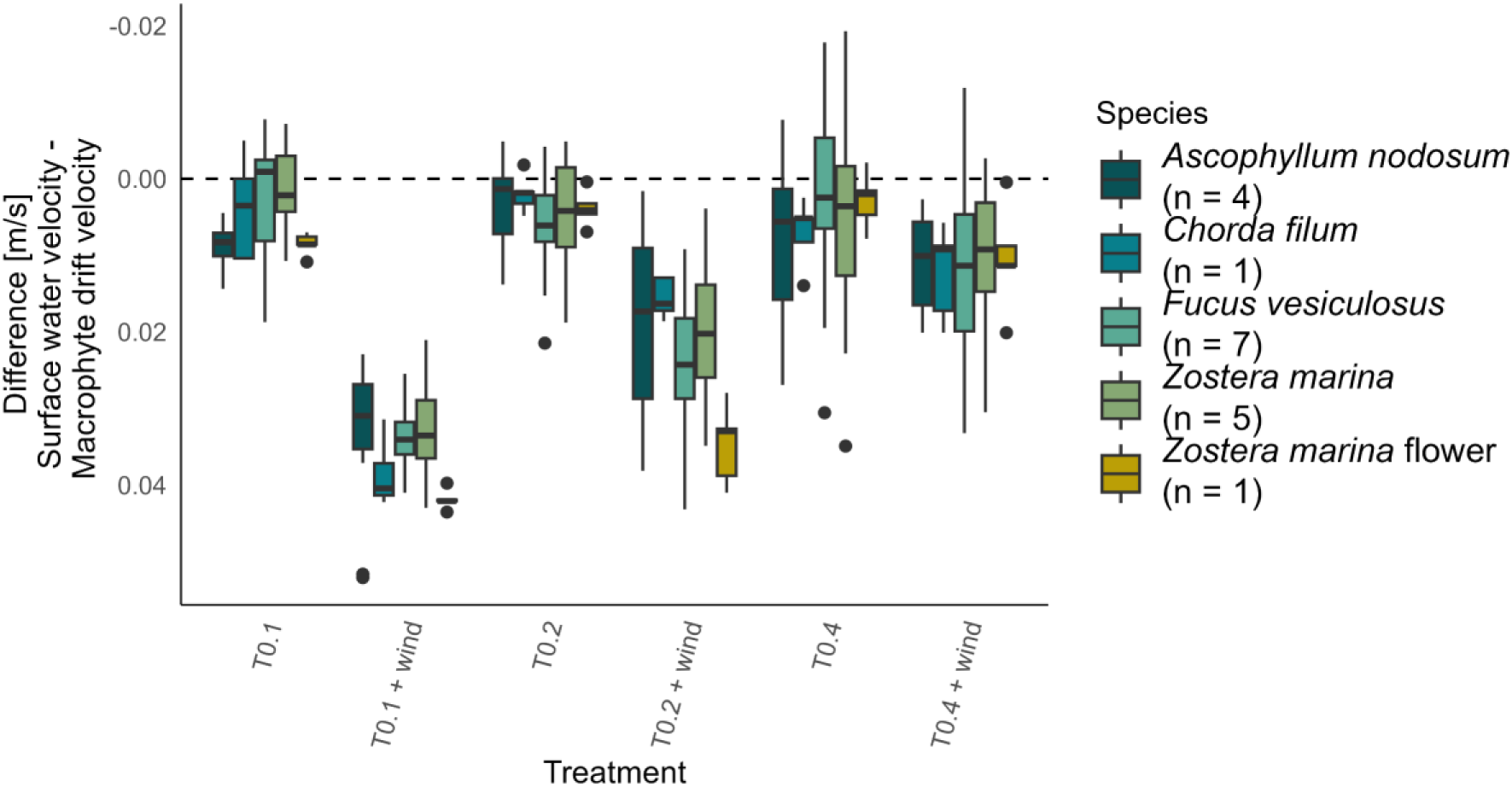
Differences between the surface water velocity and the drift velocity of the positively buoyant macrophytes. n indicates the number of individuals measured per treatment, each individual was measured five times per treatment.

Validating an existing positively buoyant species drift velocity model with and without wind The calculation of the drift velocity with the Maxey-Riley equation for floating species is based on α which is dependent on the buoyancy (δ) of a sample. The range of α is between 0.0095 and 0.0199. A clear non-linear dependence between α and δ can be seen (Figure 5), this was tested with a Spearman’s rank correlation test and revealed a perfect negative monotonic relationship (ρ = -1, p < 0.0001), indicating that as δ increases, α consistently decreases.

**Figure 5.**
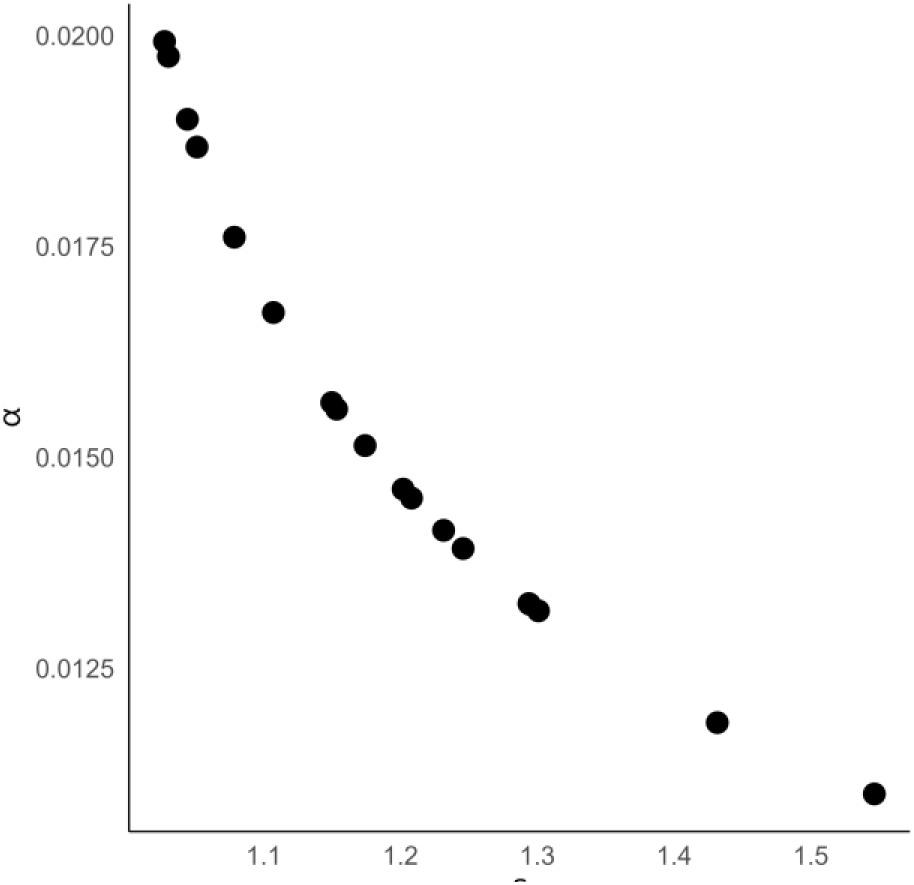
Correlation between α (buoyancy-dependent leeway factor) and δ (buoyancy) parameters of the Maxey-Riley equation. For the positively buoyant species.

For the positively buoyant species, the best model fit based on the Maxey-Riley set of equations was obtained in the treatments without wind (T0.1, T0.2, and T0.4, Supplementary Figure 9, yellow points; MSD = 0.000018 m²/s², Pearsońs *r* = 0.999, t(70) = 102.4, p < 0.001). All Pearson correlation significances were confirmed with Spearman’s correlations (Supplementary Table 3). If wind was included, method 2 (using surface water velocity with wind but without additional wind velocity) provided the best prediction of the observed drift velocities (MSD = 0.00033 m²/s², Pearsońs *r* = 0.999, t(52) = 152.61, p < 0.001, Figure 6 blue dots), this was especially visible at the highest water velocity (Figure 6C). This was followed by method 3 (combining the surface water velocity from the no-wind treatments with the added wind velocity; MSD = 0.00038 m²/s², Pearsońs *r* = 0.998, t(52) = 102.83, p < 0.001, Figure 6 dark blue triangles), which had visibly the best fit at the slowest water velocity (Figure 6A). Subsequently, the simplified leeway factor method followed (MSD = 0.00164 m²/s², Pearsońs *r* = 0.999, t(52) = 148.74, p < 0.001, Figure 6 green diamonds). The poorest fit was obtained with method 1 (incorporating both the water velocity with wind and the wind velocity; MSD = 0.00213 m²/s², Pearsońs *r* = 0.998, t(52) = 105.78, p < 0.001, Figure 6 yellow squares).

**Figure 6.**
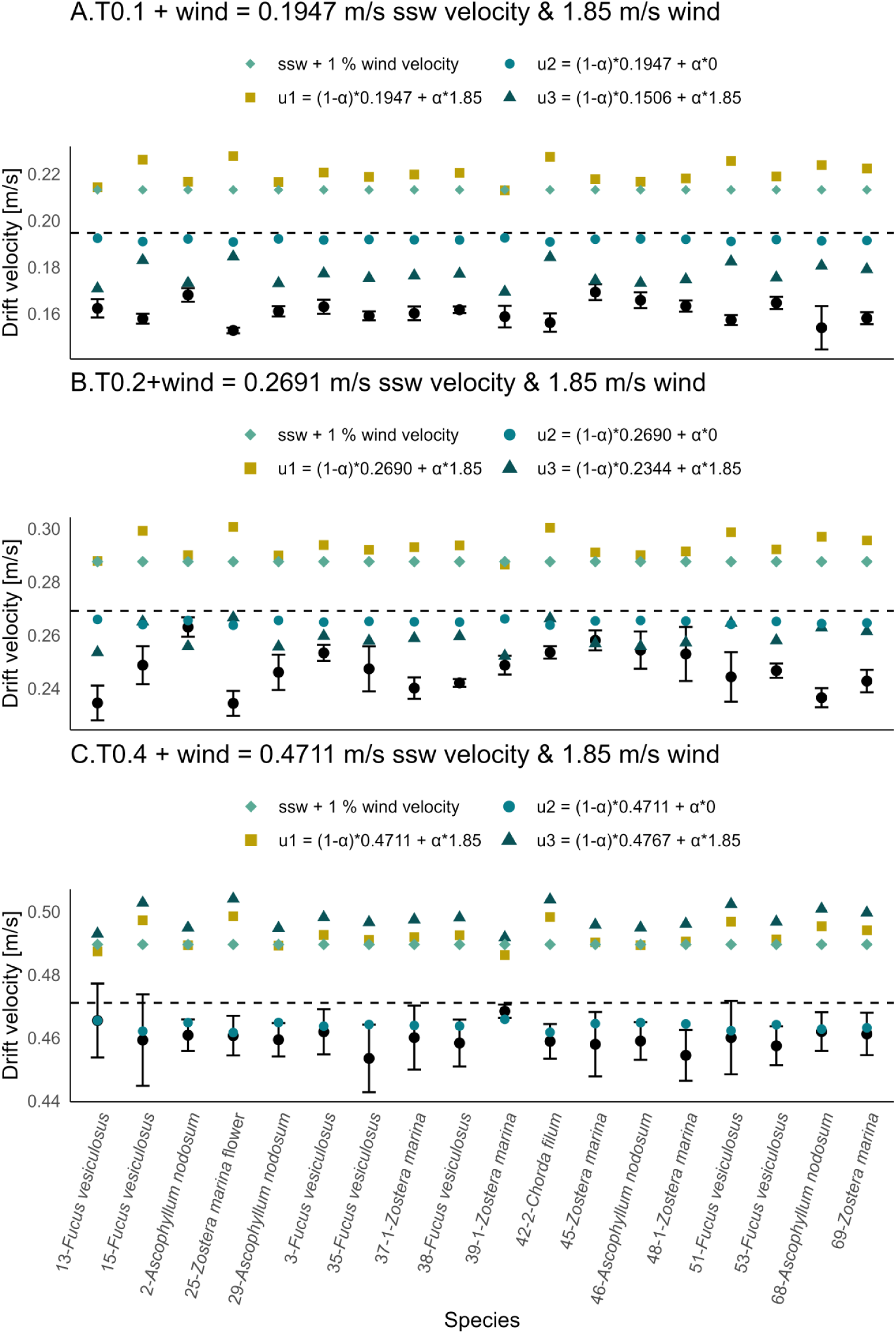
Accuracy of the Maxey-Riley model fit for each treatment with wind. The black points represent the mean ± sd of the observed drift velocities of positively buoyant species. The coloured shapes represent the drift velocities modelled using the simplified leeway factor method (green diamonds), as well as calculations with the Maxey-Riley set of equations using three different methods. Method 1 = yellow squares, Method 2 = blue points, Method 3 = dark blue triangles, for method clarification see Materials & Methods part. The dashed line indicates the ssw – sea surface water velocity.

### Model performance for macroplastic particles

As all of the macroplastic particles were positively buoyant their drift velocities were calculated with the Maxey-Riley equations. Using these equations, the calculated drift velocities of the macroplastic particles closely matched the observed values for all treatments without wind together (MSD = 0.00010 m²/s², Pearsońs *r* = 0.993, *t*(13) = 31.278, *p* < 0.001; Figure 7A – C yellow points). For the wind treatments (Figure 7D – F), the best model fit was obtained using method 2, which applies the surface water velocity from the wind treatment without adding additional wind velocity (MSD = 0.00109 m²/s², Pearsońs *r* = 0.954, *t*(13) = 11.525, *p* < 0.001, Figure 7 blue points). This was followed by method 3, which uses the surface water velocity without wind and adds the wind velocity (MSD = 0.00143 m²/s², Pearsońs *r* = 0.944, *t*(13) = 10.306, *p* < 0.001, Figure 7 dark blue triangles). The next best performance was achieved by the simplified leeway factor method, which adds 1 % of the wind velocity (MSD = 0.00190 m²/s², Pearsońs *r* = 0.953, *t*(13) = 11.322, *p* < 0.001, Figure 7 green diamonds), and the poorest fit occurred with method 1, which follows the Maxey-Riley equation as intended (MSD = 0.00202 m²/s², Pearsońs *r* = 0.942, *t*(13) = 10.15, *p* < 0.001, Figure 7 yellow squares). Also, notable is that the model was able to predict the drift velocity for P4-Juice pouch best regardless of treatment, which was the sample with the highest part emerged above the water column and nearly no submerged part. The model fit for the macroplastic particles was slightly less accurate than that for the macrophytes.

The use of the surface water velocity alone as drift velocity did not result in a better fit than the best of the tested models, both for macrophytes (MSD = 0,000022 m²/s², Pearsońs *r* = 0.999, t(55) = 183.41, p < 0.001), and macroplastic particles (MSD = 0.00028 m²/s², Pearsońs *r* = 0.993, *t*(13) = 30.624, *p* < 0.001) in the treatments without wind, as well as for macrophytes (MSD = 0.00049 m²/s², Pearsońs *r* = 0.999, t(52) = 148.74, p < 0.001), and macroplastic particles (MSD = 0.00125 m²/s², Pearsońs *r* = 0.952, *t*(13) = 11.243, *p* < 0.001) in the treatments with wind.

**Figure 7.**
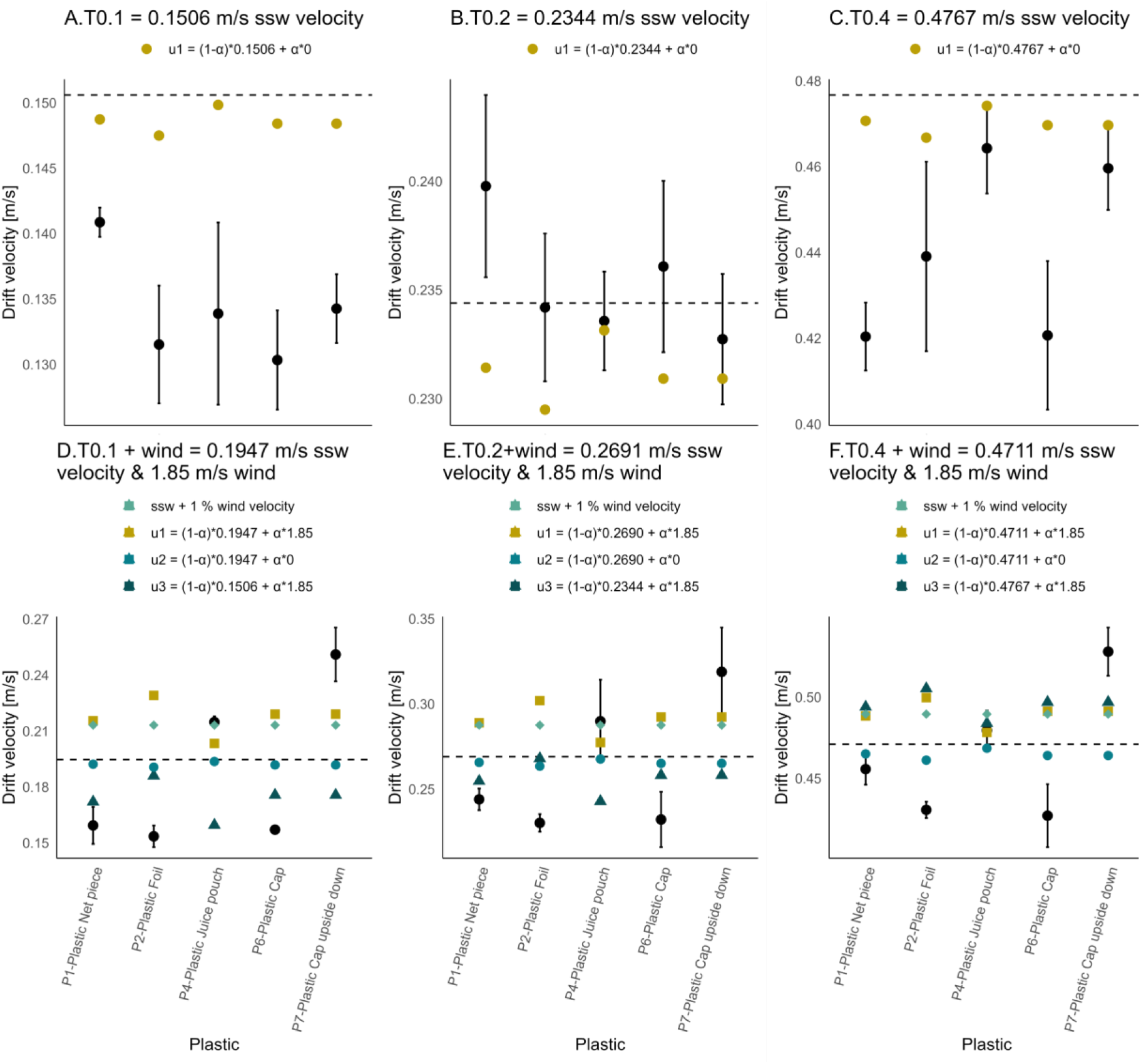
Accuracy of the Maxey-Riley model in predicting the drift of macroplastic particles. Shown for each water velocity without wind (A – C) and with wind (D – F). The black dots represent the mean ± sd of the observed drift velocities from positively buoyant species. The coloured shapes represent the drift velocities modelled using the Maxey-Riley set of equations with the three different methods. Method 1 = yellow points (A – C) or yellow squares (D – F), Method 2 = blue points, Method 3 = dark blue triangles, for method clarification see Materials & Methods part. The dashed line indicates the ssw – sea surface water velocity.

### Negatively buoyant species drift velocity model

To be able to calculate the drift velocity of negatively buoyant species, Equation 2 was developed based on morphological differences among macrophytes and their interaction with the bottom. Calibrating the bottom friction factor formulation (*BFC_Water_*) for water after Dong et al. (2023) for macrophytes yielded two scaling parameters a = 114.1662 and b = – 0.1638952. This resulted in Equation 5 to calculate the bottom friction for macrophytes

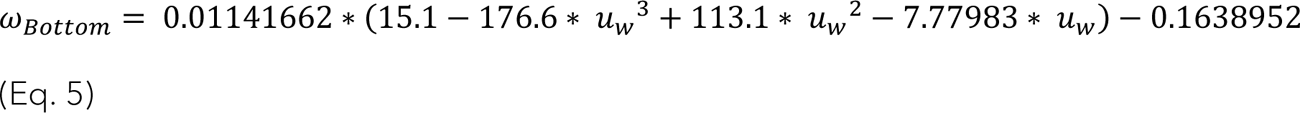

Implementing the newly developed calculation for 𝜔_𝐵𝑜𝑡𝑡𝑜𝑚_ in Equation 2, we were able to successfully predict the drift velocity of macrophytes across all three treatments without wind. The best model fit was achieved when observed terminal sedimentation velocity was used as macrophyte drag (MSD = 0.00009 m²/s², Pearsońs *r* = 0.98, *t*(56) = 42.313, *p* < 0.001, Figure 8).

Supplementary Figure 10 provides a detailed depiction of the model fit for each sample and shows that some samples would be underpredicted by the model (light blue points + error bars: mean ± sd). These correspond to negatively buoyant samples that were drifting in the water column rather than in contact with the bottom (none in T0.1, 16-2-*Spermothamnion repens*, 14-*Spermothamnion repens*, and 21-1-*Ulva clathrata* in T0.2, as well as 26-2*-Delesseria sanguinea* and 34-*Cladophora flexuosa* in T0.4). Excluding the bottom friction (𝜔_𝐵𝑜𝑡𝑡𝑜𝑚_) from the model resulted in a better fit for species drifting in the water column (𝑢 = *u_w_* − 𝜔_𝑀𝑎𝑐𝑟𝑜𝑝ℎ𝑦𝑡𝑒_ − 𝜔_𝐵𝑜𝑡𝑡𝑜𝑚_; MSD = 0.00200 m²/s², Pearsońs *r* = 0.98, *t*(3) = 9.6606, *p* < 0.05; whereas 𝑢 = *u_w_* − 𝜔_𝑀𝑎𝑐𝑟𝑜𝑝ℎ𝑦𝑡𝑒_; MSD = 0.00088 m²/s², Pearsońs *r* = 0.98, *t*(3) = 9.2078, *p* < 0.05). Interestingly, the best fit for water column species was achieved when the water velocity at the highest drift height of the macrophytes was used as their drift velocity (𝑢 = *u_w_*; MSD = 0.00046, Pearsońs *r* = 0.99, *t*(3) = 10.059, *p* < 0.05). This pattern did not hold for species drifting in contact with the bottom, for which using water velocity alone as drift velocity resulted in a poorer performance than the developed model (MSD = 0.00253 m²/s², Pearsońs *r* = 0.96, *t*(61) = 25.964, *p* < 0.001). It is also noteworthy that different samples were suspended in the water column at different water velocities.

**Figure 8.**
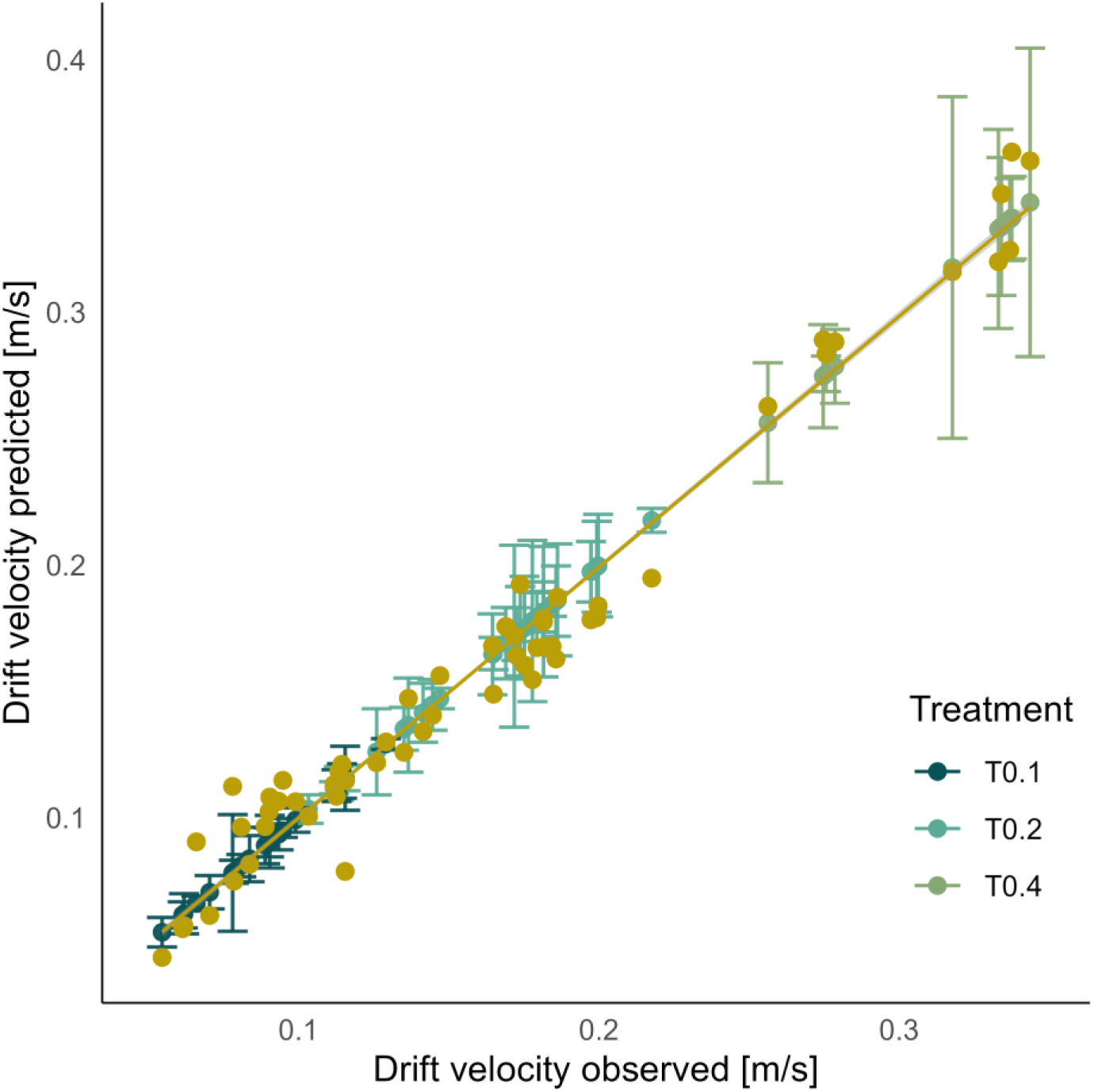
Fit of the drift velocity model for negatively buoyant species, Equation 2. The observed drift velocities in blue and green (means ± sd) vs. the modelled drift velocities in yellow or orange (means), using the observed sedimentation velocity as 𝜔_𝑀𝑎𝑐𝑟𝑜𝑝ℎ𝑦𝑡𝑒_.

## Discussion

The observed variation in drift velocities across different macrophyte and macroplastic samples suggests that characteristics unique to each sample influence drift velocity, as they would all exhibit the same drift velocity if it were solely determined by external hydrodynamic factors. These sample-specific aspects may be related to morphology or density. However, no clear trend could be identified linking specific morphological features to differences in drift velocity, despite the apparent influence of morphology on drag. The density effect arises because macrophytes and macroplastic particles are exposed to different water velocities depending on their buoyancy, ranging from the faster surface water velocity to the slower bottom water velocity. However, within the positively and negatively buoyant samples there was no clear trend between density and drift velocity, highlighting the probability of complex interactions of morphology and density. These results also emphasize that positively and negatively buoyant samples should be regarded separately, and that general statements about drifting macrophytes or macroplastic particles should be made with caution.

### The influence of wind

Wind-driven increases in surface water velocity strongly affect macrophyte drift, generally leading to higher drift velocities when wind acts in the same direction as the current. However, if wind had solely acted to increase macrophyte drift velocity, drift velocities would be expected to scale uniformly with surface water velocity. The larger discrepancy observed between surface water velocity and macrophyte drift velocity under wind forcing indicates that this is not the case. This suggests that wind primarily influences macrophyte drift velocity through changes in surface water velocity, rather than through direct aerodynamic forcing on the macrophytes themselves, while simultaneously inducing processes that decrease the drift velocity of macrophytes. Direct aerodynamic forcing on the macrophytes appears negligible, probably due to the macrophytes protruding only minimally above the water surface. This is supported by the findings of Van der Stocken et al. (2015), who showed that propagules that do not protrude above the water surface are largely unaffected by direct wind action. Coppin et al. (2024b) also reported that single kelp individuals do not present enough area above the surface to be influenced by wind. The observed reduction in drift velocity could result from wind-driven generation of small-scale turbulences, such as Langmuir circulation, a circular movement that temporarily transports particles downward (Thorpe, 2004). Support for this mechanism comes from the observed increase in vertical drift velocities at lower water velocities, indicating enhanced wind-driven turbulence in the subsurface water layers. While the macrophytes were too large to be affected by this downward transport directly, such turbulences may nevertheless have contributed to their reduced horizontal drift velocity. This effect is more pronounced the greater the difference between the surface water and the wind velocity is. In treatment 0.4 + wind, macrophyte drift velocity did not increase despite the addition of wind (in the same direction as the water velocity), coinciding with an unchanged surface water velocity, strengthening the assumption that the macrophytes are only influenced by the wind because of the effect on the surface water velocity. In contrast, the drift velocity of macroplastic particles increased even though the surface water velocity remained constant, indicating that wind had a direct effect on the macroplastic particles. This effect cannot be due to the macroplastic particles having lower densities, as their densities were in the same range as those of the macrophytes. This suggests that the observed differences are likely due to compositional variations between macroplastic particles and macrophytes, such as rigidity and surface structure.

### Macrophyte drift velocities

The dispersal of floating species typically occurs at velocities similar to those of surface water, whereas the drift velocity of species drifting on the bottom is slower (Canal-Vergés et al., 2014). This reduction in velocity, also observed in our experiments, may on the one hand reflect the decrease in water velocity with depth (Ahamed & Kundu, 2022), caused for example by friction with the seabed and energy dissipation. On the other hand, the drift velocity of bottom-drifting species is further reduced by the frictional interaction between the macrophytes and the substrate (Canal-Vergés et al., 2014). Interestingly, bottom-drifting species also exhibit greater variability in drift velocities compared to surface-drifting species. This variability is likely to reflect differences in morphological characteristics and or density between species, which influence their interactions with the substrate. As a next step, it would be interesting to examine how the macrophytes react to different bottom structures. The variability of particle drift velocities relative to the bottom water velocity varied from 17 % faster to a reduction of 58 %. Canal-Vergés et al. (2014) found a 20 % velocity reduction relative to the water velocity for green macroalgae and a 40 % reduction for brown macroalgae drifting on the bottom. In our study, there was no clear difference between green and brown macroalgae, while the highest reductions were found for red macroalgae at all three water velocities.

### Positively buoyant macrophytes and macroplastic particle drift model

Our tests have shown that the surface water velocity alone is not a sufficient predictor of the drift velocity of positively buoyant macrophytes and macroplastic particles with diverse morphologies, both with and without wind. The standard practice in the literature for incorporating wind velocity into a model is to use an ad hoc leeway factor, which is an addition of 1 – 4 % of the wind velocity to the surface water velocity, depending on the particle being modelled, from macrophytes such as *Sargassum* and *Ulva prolifera* green tides to oil slicks to shipping containers (Bonner et al., 2024; Ji et al., 2024; Jones et al., 2016; Wagner et al., 2022). At a wind velocity of 1.85 m/s, 1 % would correspond to a leeway factor of 0.0185 m/s. The set of α values (which can be seen as buoyancy-dependent leeway factors) calculated via the Maxey-Riley equation range from 0.0095 – 0.0199 (0.51 % - 1.08 % of the wind velocity), showing that an ad hoc addition of even 1 % is at the upper limit of the calculated α’s. This suggests that the simplified method of determining leeway factors would probably overestimate the effect of wind for a higher diversity of macroalgal species. It also shows that there is a noticeable difference in α values between samples. Thus, our results clearly show that it is not so easy to add a fixed value for all macrophyte species, as differences could be observed between samples. These differences depend on the buoyancy, as α and the relative density δ have a non-linear, negative monotonic relationship. It can be concluded that for a rough estimate, adding an ad hoc leeway factor can give relatively good results, but for accurate predictions of individual drift velocities it is important to calculate individual buoyancy-dependent leeway factors.

This was further supported by our comparison of model performances calculated using different methods for positively buoyant samples (macrophytes and macroplastic particles). For treatments without wind the Maxey-Riley set of equations yielded very good results. However, for treatments with wind the best results could be achieved by disregarding additional wind velocity in the Maxey-Riley equations (method 2). As mentioned above, because the macrophytes extended only slightly above the water surface, including a wind component may not be appropriate due to the limited area exposed to wind. That the models are generally slightly overpredicting the drift velocities for the majority of samples could be due to wind not only increasing surface water velocity but also inducing turbulences, which slow down the macrophytes and macroplastic particles. This would lead to the conclusion that instead of adding an additional wind velocity a factor should be deducted to account for the counter intuitive loss in drift velocity due to turbulences produced by wind. This is also supported by the result that the model was generally better at predicting treatments without wind than with wind. Another aspect that should be taken into account, is that most wind models are developed for wind speeds at 10 m above the water surface (Ji et al., 2024; Putman et al., 2018; Wagner et al., 2022). However, because wind velocity decreases by 15 – 16 % when measured at 10 m in comparison to 3 m above the sea surface (Bagiorgas et al., 2012), the actual samples experience even lower wind velocity than the models predict as they do not protrude that far above the water surface. Our study, on the other hand, better represents the effect of wind on the samples as the wind velocity measurements were taken directly at the surface, representing the wind velocities the macrophytes are truly exposed to. Using the Maxey-Riley set of equations already works very well for a majority of samples. As it is mainly dependent on buoyancy (which depends on density) and does not consider other shape parameters, in a next step, shape descriptors such as the area oriented toward the flow, or the height above or below the water column could be integrated to achieve even more differentiated results.

### Negatively buoyant macrophyte drift model

We developed the equation for the drift velocity of negatively buoyant species based on three main considerations: 1. the actual water velocity to which the macrophytes are exposed, 2. the reduction in drift velocity due to friction with the bottom, and 3. the reduction in drift velocity due to the resistance of the macrophytes to the water velocity. The improvement of the equation when incorporating the water velocity at the highest point reached by the macrophyte in the water column highlights the crucial role of accurately representing water velocity when predicting drift velocity. This underscores the necessity of accounting for the decrease in water velocity with depth in predictive models. For negatively buoyant macrophytes, friction resulting from contact with the bottom represents an important factor that must be included in the model. Here we applied an existing bottom friction factor for water from Dong et al. (2023) and adjusted its magnitude to make it applicable to macrophytes. Dong et al. (2023) proposed four equations describing the bottom friction factor for water based on different water velocities (for *u_w_* < 0.5 𝑚/𝑠, 0.5 ≤ *u_w_* < 2.2 𝑚/𝑠, 2.2 ≤ *u_w_* < 3.2 𝑚/𝑠, and *u_w_* ≥ 3.2 𝑚/𝑠). In our experiments, water velocities reached up to ∼ 0.5 m/s, allowing us to evaluate the applicability of the shape of the first equation for macrophyte bottom friction. Future studies with higher water velocities would allow us to evaluate the model’s performance under a broader range of water velocities. It should be noted that the experiments were conducted on a smooth PVC bottom, which may underestimate the role of bottom friction in natural environments. As a next step, more diverse bottom structures could be tested to better reflect natural conditions. The resistance of macrophytes to water velocity was determined experimentally from their sedimentation velocities. However, in cases where such measurements are unavailable, calculating macrophyte drag using the sedimentation velocity equation proposed by Gronwald et al. (2025) also achieved a very good fit (MSD = 0.00013 m²/s², Pearsońs *r* = 0.97, *t*(56) = 30.217, *p* < 0.001, Supplementary Figure 11). A visual comparison of the results obtained using the observed versus the calculated sedimentation velocities is shown in Supplementary Figure 11. This approach enables the calculation of drift velocities for negatively buoyant species without the need for experimental measurements. So far, the individual shape of different macrophyte species was, to our knowledge, disregarded in models predicting drift velocity. This is incorporated now by building on the sedimentation velocity model, which already includes a shape factor (Gronwald et al., 2025). Together, these three aspects enable accurate predictions of drift velocity for negatively buoyant macrophytes based on their morphology, thereby filling a critical gap in particle transport modelling. Improved predictions of macrophyte drift can enhance our ability to identify where drifting macrophytes may affect an ecosystem, for example by reaching new habitats and facilitating colonization. Such processes may be particularly important under climate change scenarios, as they could support species persistence if original habitats become unsuitable. In addition, this approach provides a valuable tool for assessing population connectivity and predicting potential beaching locations.

Between the samples drifting at the surface and those drifting along the bottom were samples that were negatively buoyant but not in contact with the bottom, instead drifting within the water column. Unlike positively and negatively buoyant samples, the drift velocity of these intermediate samples was best predicted by the surrounding water velocity, suggesting that they are not substantially influenced by bottom friction or drag dependent on density and shape. This highlights the importance of a macrophyte’s position within the water column for accurately predicting drift velocity and, consequently, its dispersal.

## Conclusion

In conclusion, the drift velocity of macrophytes depends on their position in the water column and therefore on the water velocity, which is shaped by hydrodynamic factors. While wind increases the drift velocity of macrophytes by increasing the surface water velocity, its full impact is counteracted to some degree by the turbulence it induces, which reduces the drift velocity. Water velocity alone was insufficient at predicting the drift velocity of macrophytes for species drifting on the surface or on the bottom, highlighting the need for more complex models. For positively buoyant samples, existing buoyancy-based models are already quite good at predicting drift velocity of particles such as macrophytes or macroplastic debris when aerodynamic particle drag is not taken into consideration in addition to the wind-increased surface water velocity. Interestingly, for positively buoyant samples, the methods that best predicted macrophyte drift velocity also performed best for macroplastic particles, highlighting their generality. Negatively buoyant species show more variable drift velocities, probably due to their morphologically different interactions with the bottom. To our knowledge, this is the first time that an equation has been developed to calculate the drift velocity of negatively buoyant macrophytes, taking into account their different shapes and interactions with the bottom, as well as macrophyte drag.

Data will be made available on PANGEA once publication is accepted (https://doi.pangaea.de/10.1594/PANGAEA.992527).

## Acknowledgments

We thank Johanna Behrisch for creating the illustration used in the graphical abstract.

## Credit Statement

FG, FW and TB initiated and designed this study. FG, FW, ZZ collected the data. FG analysed the data. FG generated the models. FG wrote the original draft. All authors contributed to the final version of the manuscript.

## References

Ahamed, N., & Kundu, S. (2022). Application of the fractional entropy for one-dimensional velocity distribution with dip-phenomenon in open-channel turbulent flows. Stochastic Environmental Research and Risk Assessment, 36, 1289–1312. 10.1007/s00477-022-02210-5

Arroyo, N., & Bonsdorff, E. (2016). The Role of Drifting Algae for Marine Biodiversity. In E. Ólafsson (Ed.), Marine macrophytes as foundation species (pp. 100–123). CRC Press, Taylor & Francis Group. 10.4324/9781315370781-6

Bagiorgas, H., Mihalakakou, G., & Al-Hadhrami, L. (2012). Offshore wind speed and wind power characteristics for ten locations in Aegean and Ionian Seas. Journal of Earth System Science, 121. 10.1007/s12040-012-0203-9

Beron-Vera, F. J. (2020). Nonlinear dynamics of inertial particles in the ocean: From drifters and floats to marine debris and Sargassum. 10.48550/arXiv.2007.15638

Beron-Vera, F. J. (2024). Dynamics of inertial particles on the ocean surface with unrestricted reserve buoyancy. Physics of Fluids, 36(10), 101702. 10.1063/5.0226779

Beron-Vera, F. J., Olascoaga, M. J., & Miron, P. (2019). Building a Maxey–Riley framework for surface ocean inertial particle dynamics. Physics of Fluids, 31(9), Article 9. 10.1063/1.5110731

Bhaganagar, K., Kolar, P., Faruqui, S. H. A., Bhattacharjee, D., Alaeddini, A., & Subbarao, K. (2022). A Novel Machine-Learning Framework With a Moving Platform for Maritime Drift Calculations. Frontiers in Marine Science, 9, 831501. 10.3389/fmars.2022.831501

Biber, P. D. (2007). Transport and persistence of drifting macroalgae (Rhodophyta) are strongly influenced by flow velocity and substratum complexity in tropical seagrass habitats. Marine Ecology Progress Series, 343, 115–122. 10.3354/meps06893

Björk, M., Rosenqvist, G., Gröndahl, F., & Bonaglia, S. (2023). Methane emissions from macrophyte beach wrack on Baltic seashores. Ambio, 52, 171–181. 10.1007/s13280-022-01774-4

Bonner, G., Beron-Vera, F. J., & Olascoaga, M. J. (2024). Charting the course of *Sargassum*: Incorporating nonlinear elastic interactions and life cycles in the Maxey–Riley model. PNAS Nexus, 3(10), pgae451. 10.1093/pnasnexus/pgae451

Canal-Vergés, P., Potthoff, M., Hansen, F. T., Holmboe, N., Rasmussen, E. K., & Flindt, M. R. (2014). Eelgrass re-establishment in shallow estuaries is affected by drifting macroalgae – Evaluated by agent-based modeling. Ecological Modelling, 272, 116–128. 10.1016/j.ecolmodel.2013.09.008

Christensen, A., Murawski, J., She, J., & John, M. S. (2023). Simulating transport and distribution of marine macro-plastic in the Baltic Sea. PLOS ONE, 18(1), e0280644. 10.1371/journal.pone.0280644

Coppin, R., Rautenbach, C., & Smit, A. (2024a). Individual-based numerical experiment to describe the distribution of floating kelp within the Southern Benguela Upwelling System. Botanica Marina, 67(5), 469–486. 10.1515/bot-2023-0061

Coppin, R., Rautenbach, C., & Smit, A. (2024b). Numerical experiments investigating the influence of drag on trajectory patterns of floating macroalgae. Botanica Marina, 67(5), 449–468. 10.1515/bot-2023-0059

Delpeche-Ellmann, N., Giudici, A., Rätsep, M., & Soomere, T. (2021). Observations of surface drift and effects induced by wind and surface waves in the Baltic Sea for the period 2011–2018. Estuarine, Coastal and Shelf Science, 249, 107071. 10.1016/j.ecss.2020.107071

Dong, Y., Jiang, J., Liu, X., Wang, D., & Zhang, J. (2023). An empirical formula of bottom friction coefficient with a dependence on the current speed for the tidal models. Frontiers in Marine Science, 10, 1206024. 10.3389/fmars.2023.1206024

Eriksen, M., Cowger, W., Erdle, L., Coffin, S., Villarrubia-Gómez, P., Moore, C., Carpenter, E., Day, R., Thiel, M., & Wilcox, C. (2023). A growing plastic smog, now estimated to be over 170 trillion plastic particles afloat in the world’s oceans—Urgent solutions required. PLOS ONE, 18, e0281596. 10.1371/journal.pone.0281596

Flindt, M., Neto, J. M., Amos, C., Pardal, M., Bergamasco, A., Pedersen, C., & Andersen, F. (2004). Plant bound nutrient transport. Mass transport in estuaries and lagoons. In S. L. Nielsen, G. T. Banta, & M. F. Pedersen (Eds.), The influence of primary producers on estuarine nutrient cycling: The Fate of Nutrient Biomass. (pp. 93–128). 10.1007/978-1-4020-3021-5_4

Gronwald, F., Weinberger, F., Bouma, T. J., & Karez, R. (2025). Sedimentation and drag in drifting macrophytes and plastic objects: A model. Scientific Reports, 15(1), 43088. 10.1038/s41598-025-28893-8

Holmquist, J. G. (1994). Benthic macroalgae as a dispersal mechanism for fauna: Influence of a marine tumbleweed. Journal of Experimental Marine Biology and Ecology, 180(2), 235–251. 10.1016/0022-0981(94)90069-8

Jachowski, J., & Książkiewicz, E. (2024). Numerical Prediction of Pneumatic Life Raft Performance. *TransNav*, International Journal on Marine Navigation and Safety Od Sea Transportation, 18(1), 229–232. 10.12716/1001.18.01.24

Ji, M., Dou, X., Zhao, C., & Zhu, J. (2024). Exploring the Green Tide Transport Mechanisms and Evaluating Leeway Coefficient Estimation via Moderate-Resolution Geostationary Images. Remote Sensing, 16, 2934. 10.3390/rs16162934

Jones, C. E., Dagestad, K.-F., Breivik, Ø., Holt, B., Röhrs, J., Christensen, K. H., Espeseth, M., Brekke, C., & Skrunes, S. (2016). Measurement and modeling of oil slick transport. Journal of Geophysical Research: Oceans, 121(10), 7759–7775. 10.1002/2016JC012113

Lai, S., Yaakub, S., Poh, T., Bouma, T., & Todd, P. (2018). Unlikely Nomads: Settlement, Establishment, and Dislodgement Processes of Vegetative Seagrass Fragments. Frontiers in Plant Science, 9, 160. 10.3389/fpls.2018.00160

Li, J., Wang, Z., Li, W., Jing, S., Graco-Roza, C., & Arvola, L. (2025). Impact of the Reynolds Numbers on the Velocity of Floating Microplastics in Open Channels. Water, 17(4), 588. 10.3390/w17040588

Miron, P., Medina, S., Olascoaga, M. J., & Beron-Vera, F. J. (2020). Laboratory verification of the buoyancy dependence of the carrying flow in a Maxey–Riley theory for inertial ocean dynamics. Physics of Fluids, 32(7), 071703. 10.1063/5.0018272

Novelli, G., Guigand, C., Cousin, C., Ryan, E., Laxague, N., Dai, H., Haus, B., & Özgökmen, T. (2017). A Biodegradable Surface Drifter for Ocean Sampling on a Massive Scale. Journal of Atmospheric and Oceanic Technology, 34(11), 2509–2532. 10.1175/JTECH-D-17-0055.1

Ortega, A., Geraldi, N. R., Alam, I., Kamau, A. A., Acinas, S. G., Logares, R., Gasol, J. M., Massana, R., Krause-Jensen, D., & Duarte, C. M. (2019). Important contribution of macroalgae to oceanic carbon sequestration. Nature Geoscience, 12(9), 748–754. 10.1038/s41561-019-0421-8

Putman, N., Goni, G., Gramer, L., Hu, C., Johns, E., Triñanes, J., & Wang, M. (2018). Simulating transport pathways of pelagic *Sargassum* from the Equatorial Atlantic into the Caribbean Sea. Progress in Oceanography, 165, 205–214. 10.1016/j.pocean.2018.06.009

Riazi, A., & Türker, U. (2019). The drag coefficient and settling velocity of natural sediment particles. Computational Particle Mechanics, 6(3), 427–437. 10.1007/s40571-019-00223-6

Sharqawy, M. H., Lienhard, J. H., & Zubair, S. M. (2010). Thermophysical properties of seawater: A review of existing correlations and data. Desalination and Water Treatment, 16(1), 354–380. 10.5004/dwt.2010.1079

Thorpe, S. A. (2004). LANGMUIR CIRCULATION. Annual Review of Fluid Mechanics, 36, 55–79. 10.1146/annurev.fluid.36.052203.071431

van den Hoek, C. (1987). The possible significance of long-range dispersal for the biogeography of seaweeds. Helgoländer Meeresuntersuchungen, 41(3), 261–272. 10.1007/BF02366191

Van der Stocken, T., Vanschoenwinkel, B., Ryck, D. J. R. D., Bouma, T. J., Dahdouh-Guebas, F., & Koedam, N. (2015). Interaction between Water and Wind as a Driver of Passive Dispersal in Mangroves. PLOS ONE, 10(3), e0121593. 10.1371/journal.pone.0121593

Wagner, T. J. W., Eisenman, I., Ceroli, A. M., & Constantinou, N. C. (2022). How Winds and Ocean Currents Influence the Drift of Floating Objects. Journal of Physical Oceanography, 52(5), 907–916. 10.1175/JPO-D-20-0275.1

Wu, J., Cheng, L., Chu, S., & Song, Y. (2024). An autonomous coverage path planning algorithm for maritime search and rescue of persons-in-water based on deep reinforcement learning. Ocean Engineering, 291, 116403. 10.1016/j.oceaneng.2023.116403

Zhou, F., Ge, J., Liu, D., Ding, P., Chen, C., & Wei, X. (2021). The Lagrangian-based Floating Macroalgal Growth and Drift Model (FMGDM v1.0): Application to the Yellow Sea green tide. Geoscientific Model Development, 14(10), 6049–6070. 10.5194/gmd-14-6049-2021

