## Supplementary material for "How wind and currents shape the drift velocity of macrophytes and macroplastic particles – from experiment to model"

### Supplementary tables

Supplementary Table 1. Kruskal-Wallis test results to test for significant differences within each position for each Treatment. $X^2$ - chi squared, df – degrees of freedom, significant results are indicated by *.

| | $X^2$ | Df | p-value | |
|---|---|---|---|---|
| | Treatment 0.1 | | | |
| Surface | 98.052 | 23 | < 0.001 | * |
| Bottom | 97.663 | 22 | < 0.001 | * |
| | Treatment 0.2 | | | |
| Surface | 68.83 | 30 | < 0.001 | * |
| Bottom | 133.93 | 35 | < 0.001 | * |
| | Treatment 0.4 | | | |
| Surface | 69.94 | 23 | < 0.001 | * |
| Bottom | 41.648 | 14 | < 0.001 | * |

Supplementary Table 2. Testing for the divergence of the drift velocity from the surface water velocity. If the difference in surface water velocity – drift velocity was normally distributed a one-sample t-test was used, if the data were not normally distributed a Wilcoxon signed-rank test was used.

| Treatment | Test | Test statistic (V) | t-value | df | p-value | 95 % Confidence interval | Sample mean |
|---|---|---|---|---|---|---|---|
| T0.1 | Wilcoxon signed-rank test | 3350 | - | - | < 0.001 | - | - |
| T0.1 + wind | One-sample t-test | - | 57.31 | 90 | < 0.001 | [0.0328, 0.0351] | 0.0340 |
| T0.2 | One-sample t-test | - | 8.2332 | 90 | < 0.001 | [0.0035, 0.0058] | 0.0047 |
| T0.2 + wind | One-sample t-test | - | 21.877 | 90 | < 0.001 | [0.0200, 0.0240] | 0.0220 |
| T0.4 | Wilcoxon signed-rank test | 2986 | - | - | < 0.001 | - | - |
| T0.4 + wind | One-sample t-test | - | 11.815 | 90 | < 0.001 | [0.0093, 0.0130] | 0.0112 |

Supplementary Table 3. Spearman´s rank correlation coefficient (ρ) and p-values for all models in which the data were not normally distributed.

| Treatment | Spearman`s ρ | p-value |
|---|---|---|
| **Positively buoyant species** | | |
| T0.1, T0.2, T0.4 | 0.87 | < 0.001 |
| Water velocity without wind | 0.94 | < 0.001 |
| Method 1 | 0.85 | < 0.001 |
| Method 2 | 0.93 | < 0.001 |
| Method 3 | 0.85 | < 0.001 |
| Simplified leeway factor method | 0.94 | < 0.001 |
| Water velocity with wind | 0.94 | < 0.001 |
| **Macroplastic particles** | | |
| T0.1, T0.2, T0.4 | 0.91 | < 0.001 |
| Water velocity without wind | 0.94 | < 0.001 |
| Method 1 | 0.81 | < 0.001 |
| Method 2 | 0.87 | < 0.001 |
| Method 3 | 0.81 | < 0.001 |
| Simplified leeway factor method | 0.89 | < 0.001 |
| Water velocity with wind | 0.89 | < 0.001 |
| **Negatively buoyant species** | | |
| Observed drag | 0.97 | < 0.001 |
| Calculated drag | 0.94 | < 0.001 |
| Only water velocity | 0.93 | < 0.001 |

### Supplementary figures

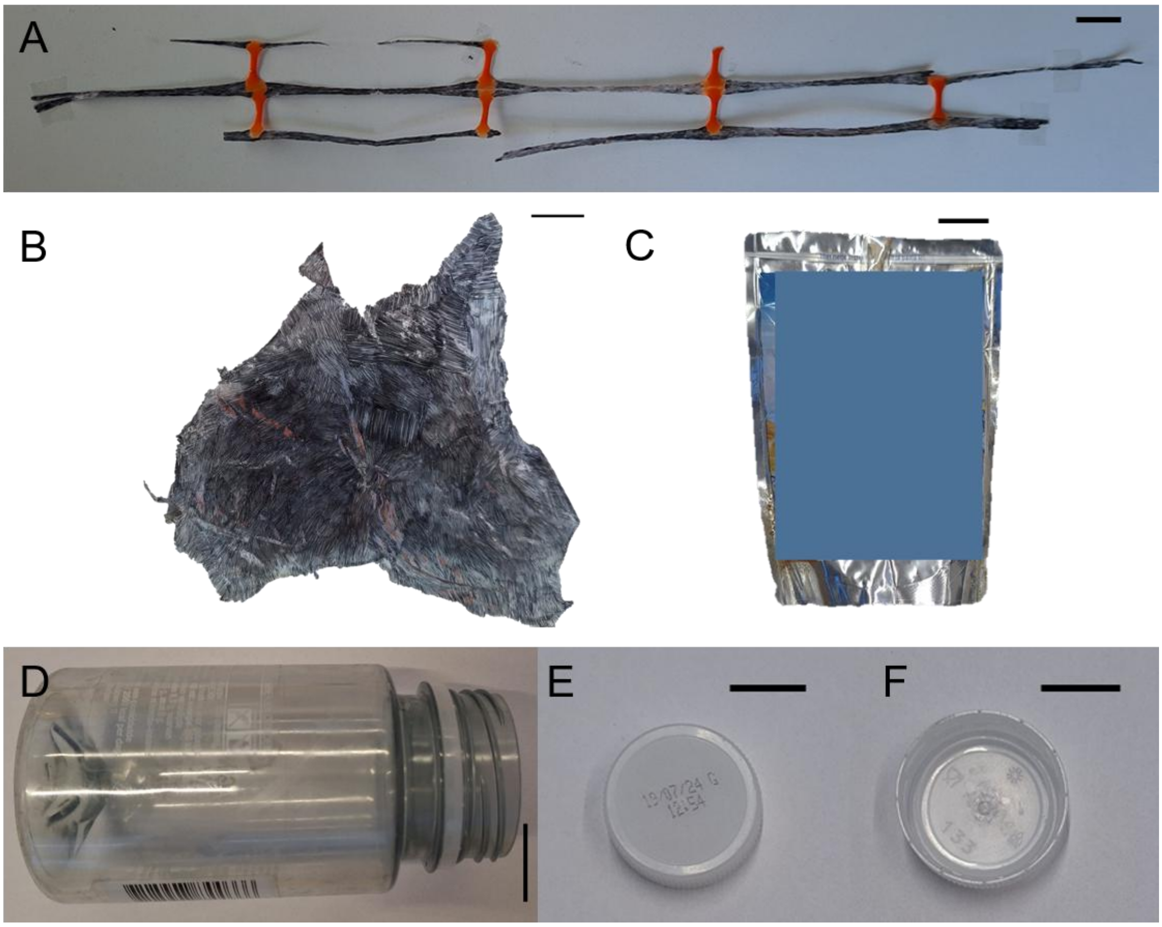


Supplementary Figure 1. Macroplastic particles. (A) P1-Net piece, (B) P2-Foil, (C) P4-Juice pouch, (D) P5-Bottle, (E) P6-Cap, (F) P7-Cap upside down. Each scale bar corresponds to 2 cm.

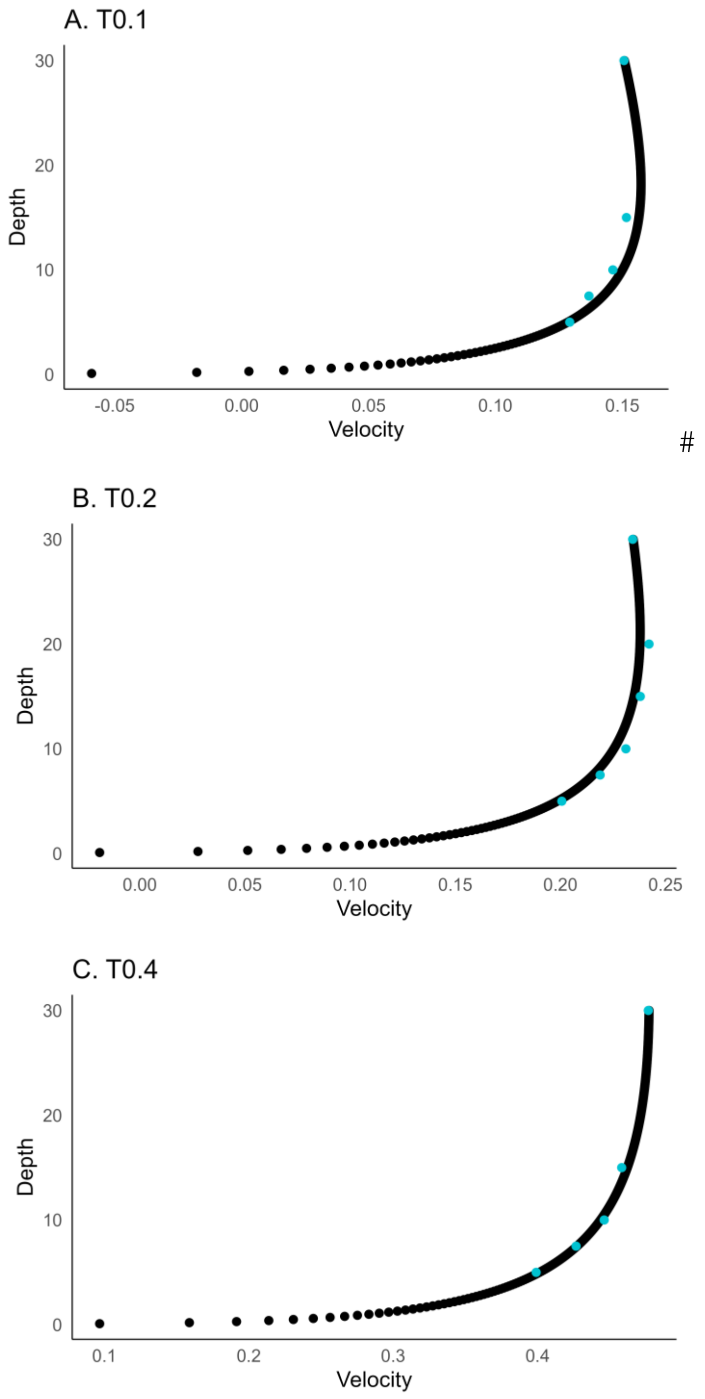


Supplementary Figure 2. Fit of the modelled water velocity dependent on depth equation (Eq. 3) for treatments T0.1 (A), T0.2 (B), and T0.4 (C). Black dots show the fitted values from the model, and blue dots the measured water velocities.

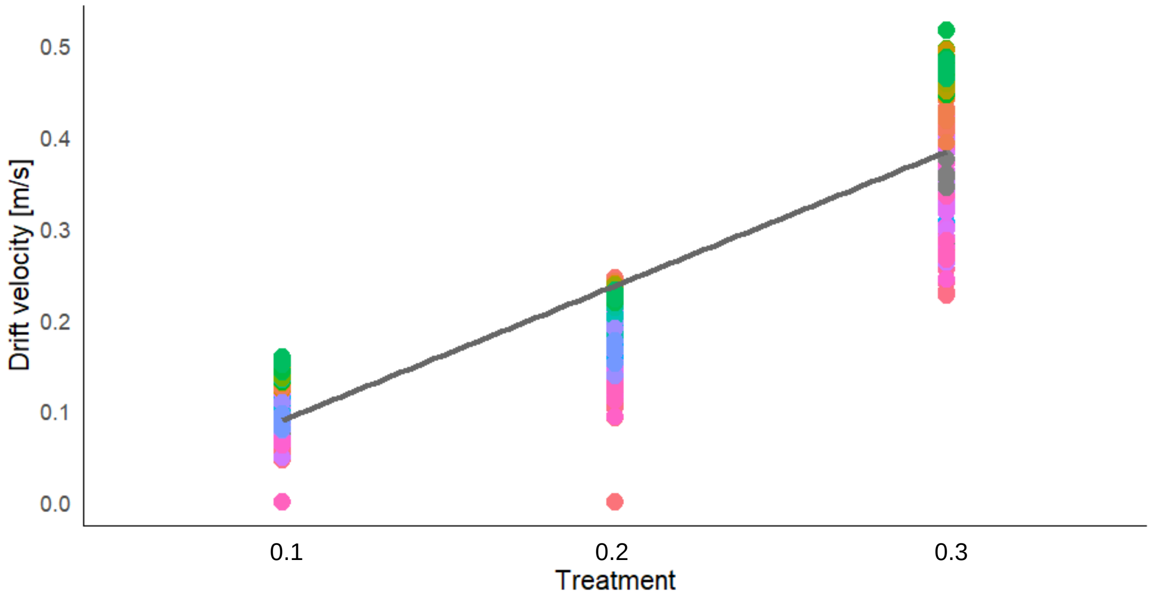


Supplementary Figure 3. Linear correlation between the drift velocities of the samples at each treatment. Colours are representing samples.

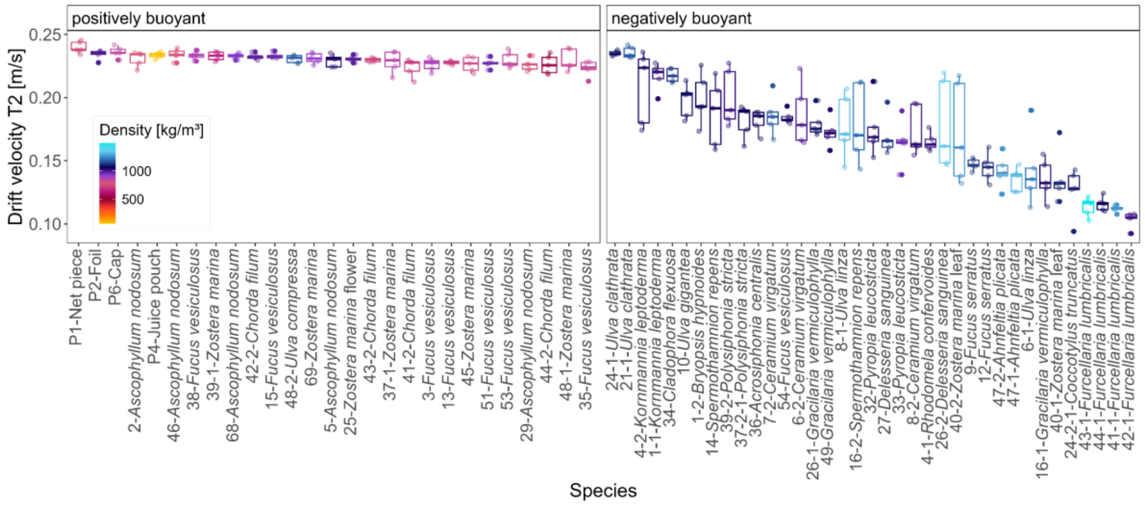


Supplementary Figure 4. Positively (left panel) and negatively (right panel) buoyant species coloured according to their density and sorted by their drift velocity in treatment 2.

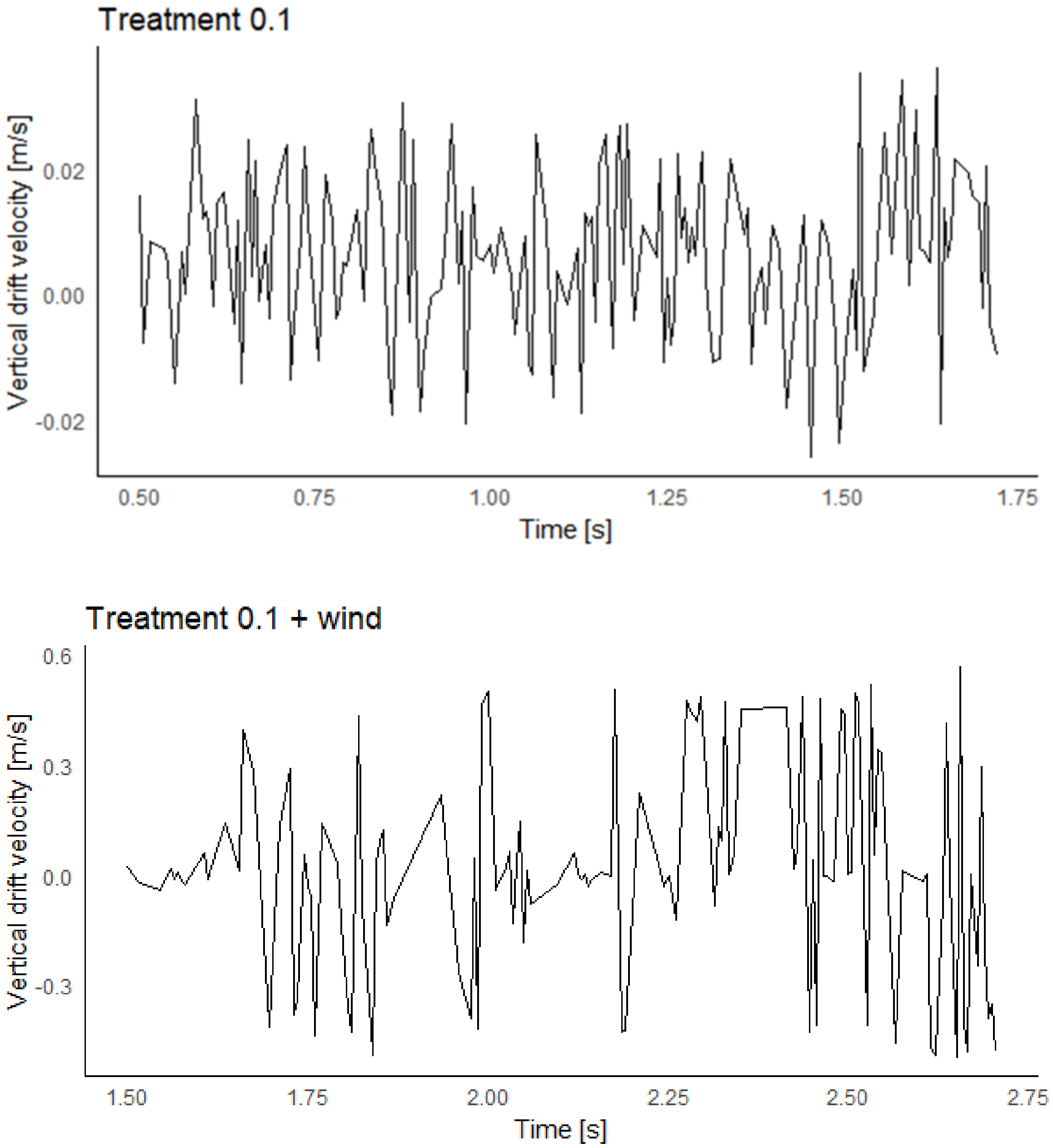


Supplementary Figure 5. Vertical drift velocity of water particles at 15 cm water height from the bottom, for Treatment 0.1 with and without wind. Vertical drift velocities were despiked.

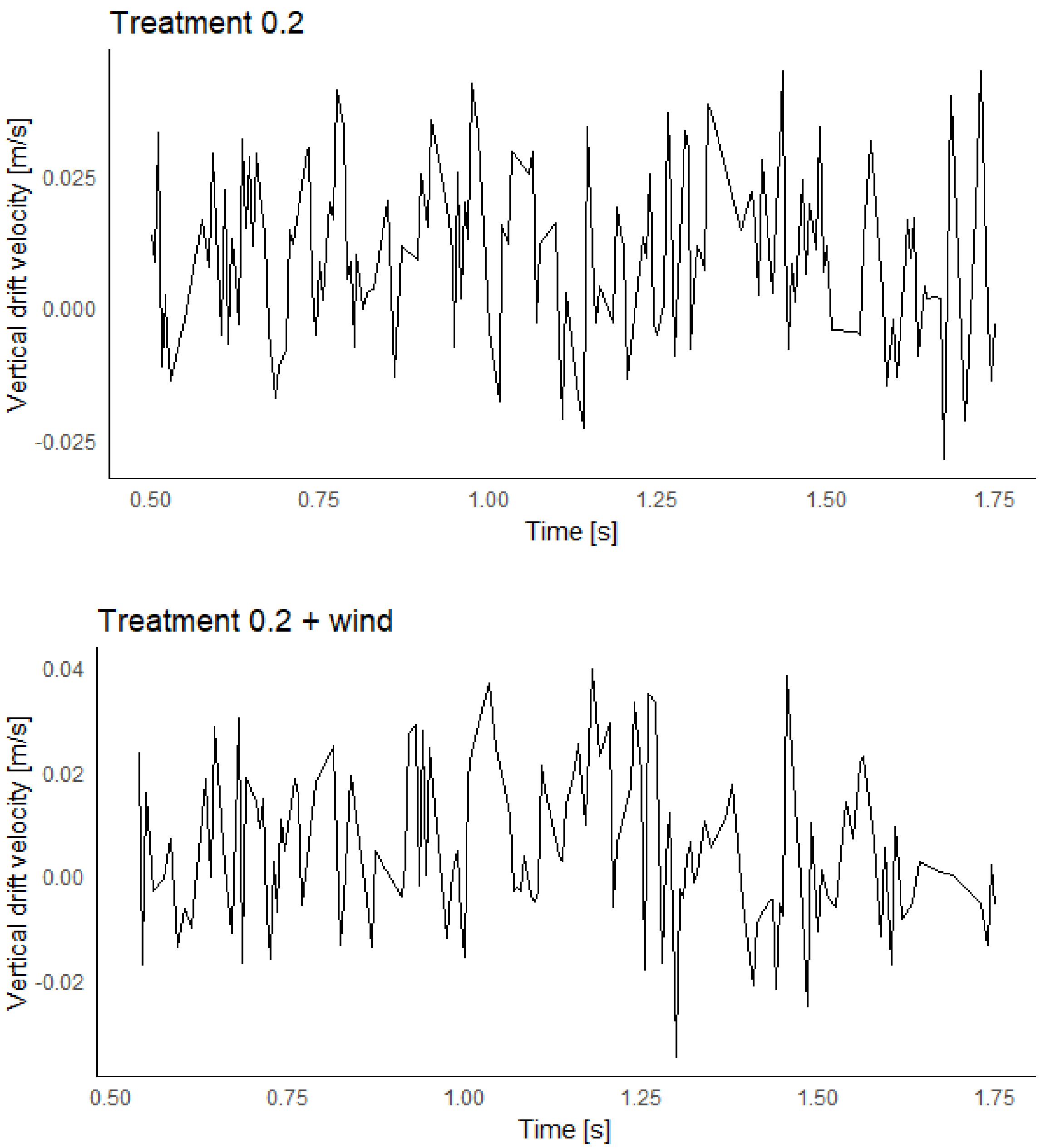


Supplementary Figure 6. Vertical drift velocity of water particles at 15 cm water height from the bottom, for Treatment 0.2 with and without wind. Vertical drift velocities were despiked.

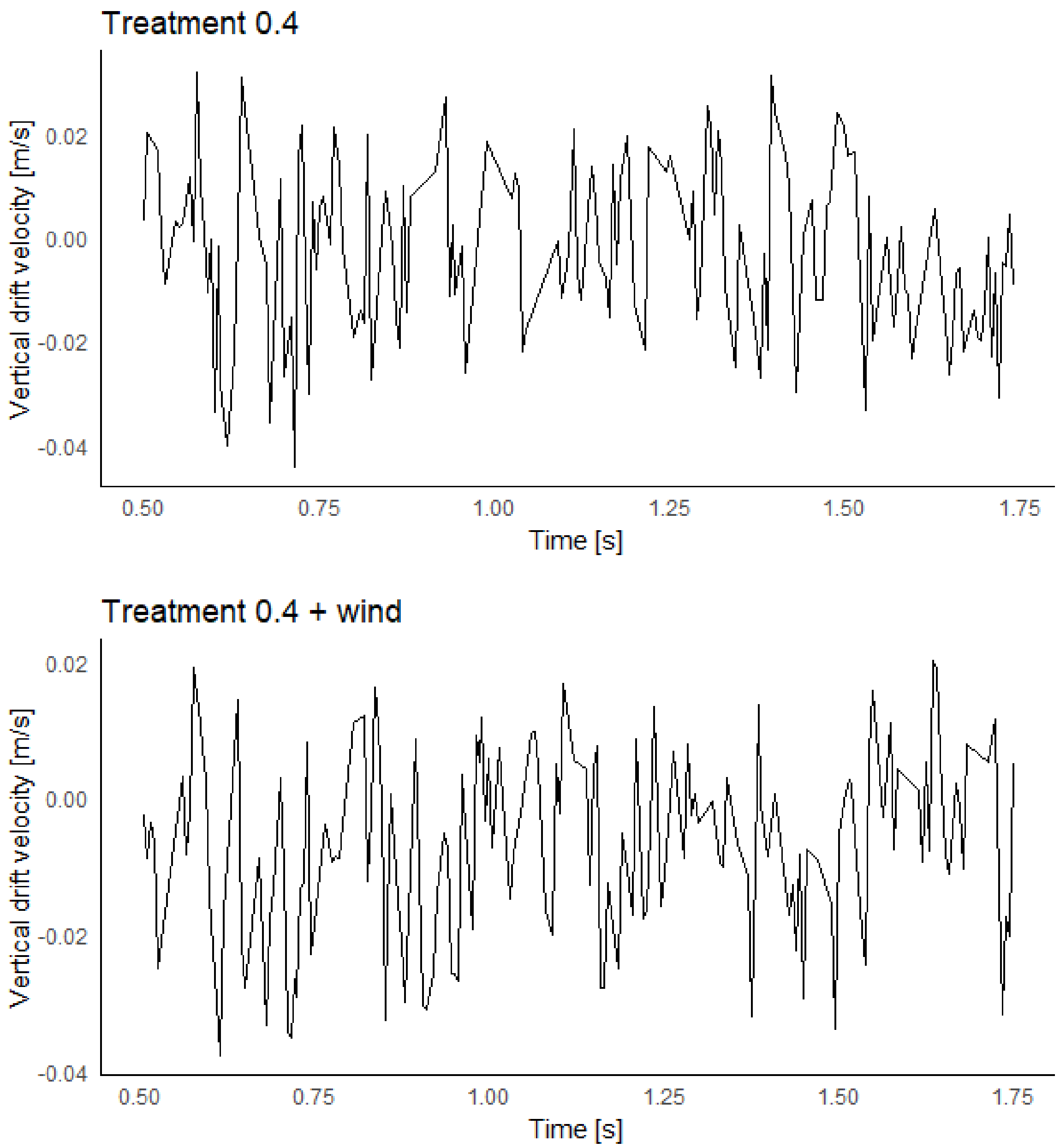


Supplementary Figure 7. Vertical drift velocity of water particles at 15 cm water height from the bottom, for Treatment 0.4 with and without wind. Vertical drift velocities were despiked.

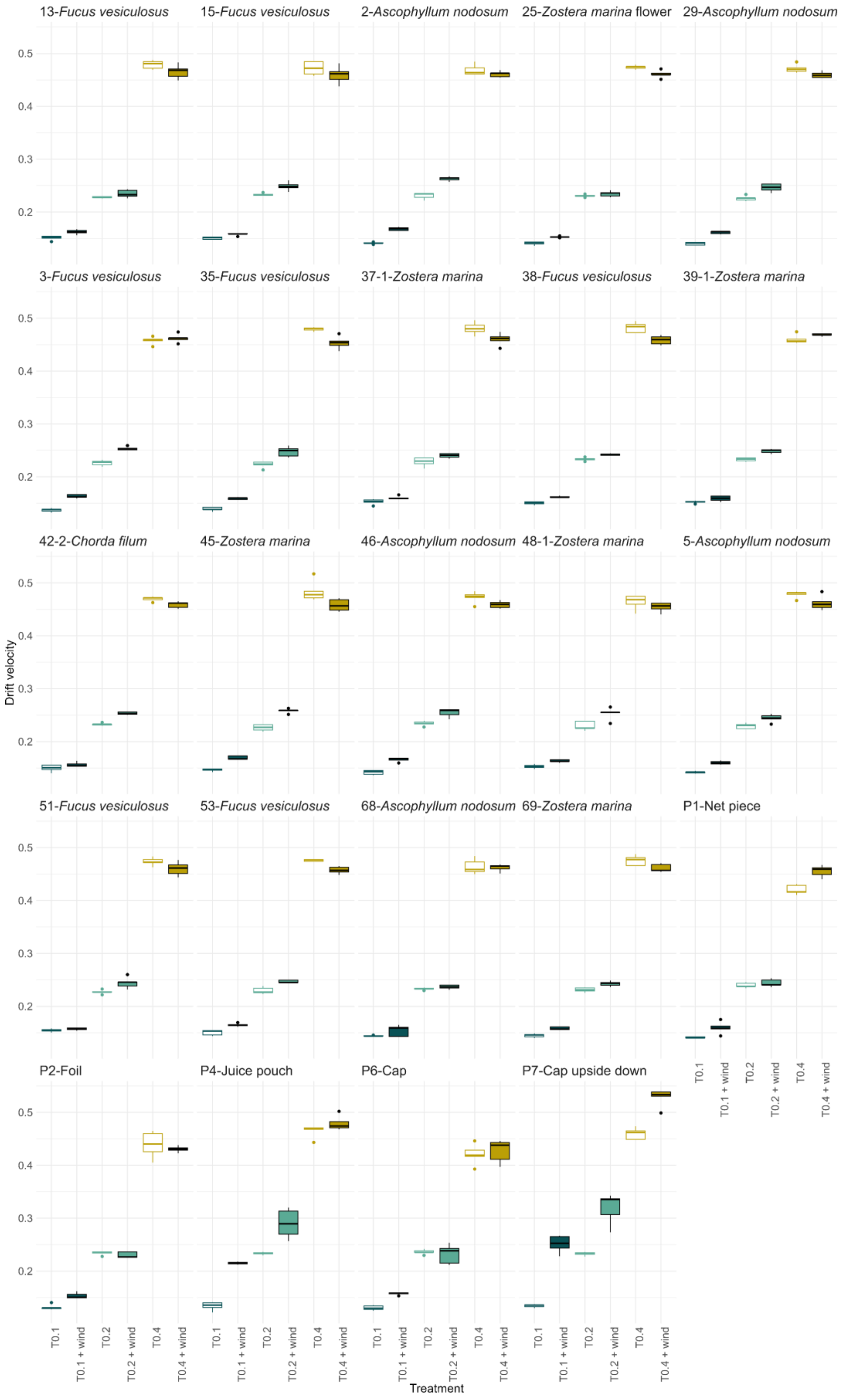


Supplementary Figure 8. Drift velocities for each positively buoyant macrophyte and macroplastic particle individual for the three current velocities with and without wind.

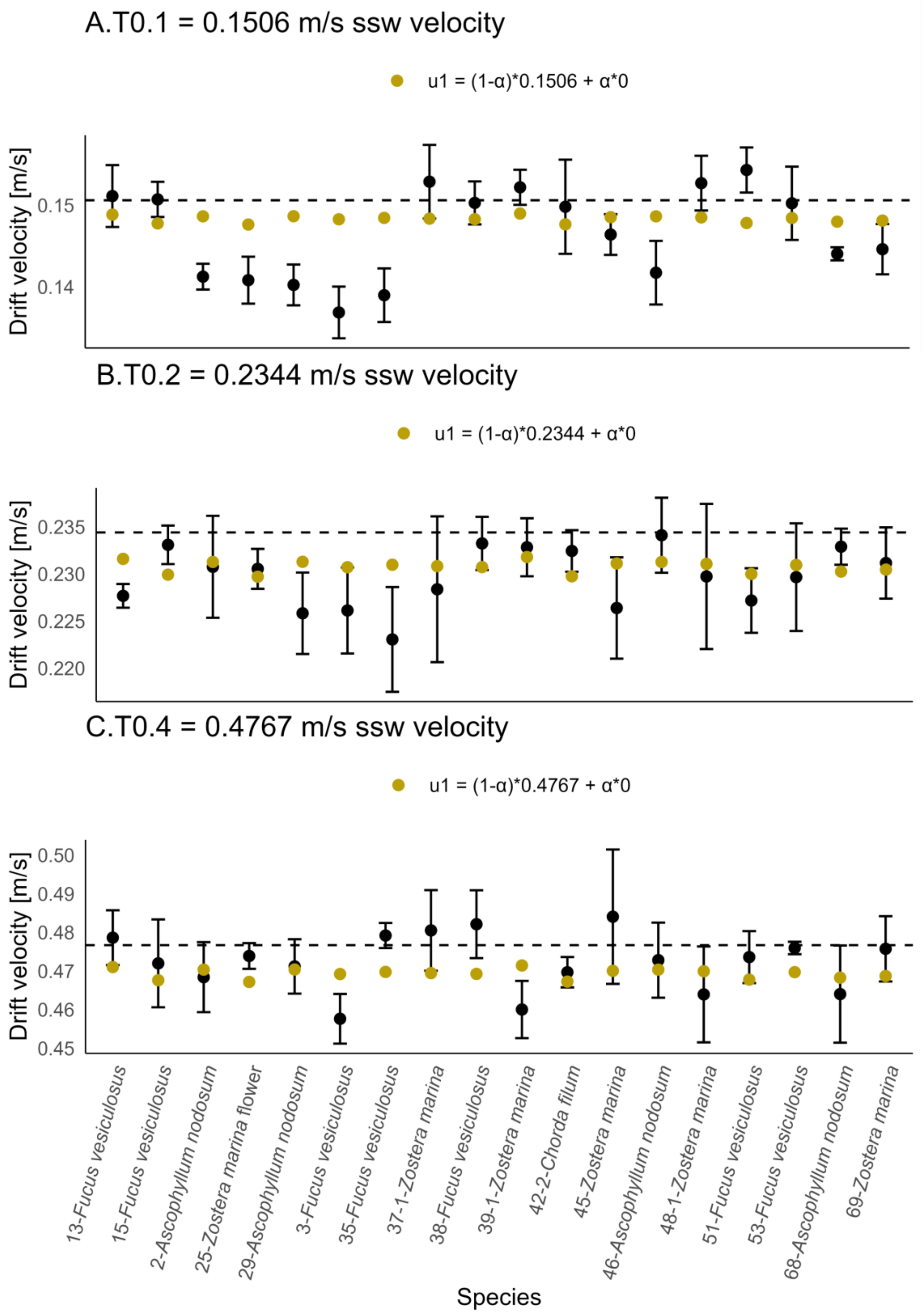


Supplementary Figure 9. Accuracy of the Maxey-Riley model fit for each treatment without wind. The black points represent the mean ± sd of the observed drift velocities of positively buoyant species. The green points represent the drift velocities modelled using method 1 of the Maxey-Riley set of equations, see Materials & Methods for method explanation. The dashed line indicates the ssw – sea surface water velocity.

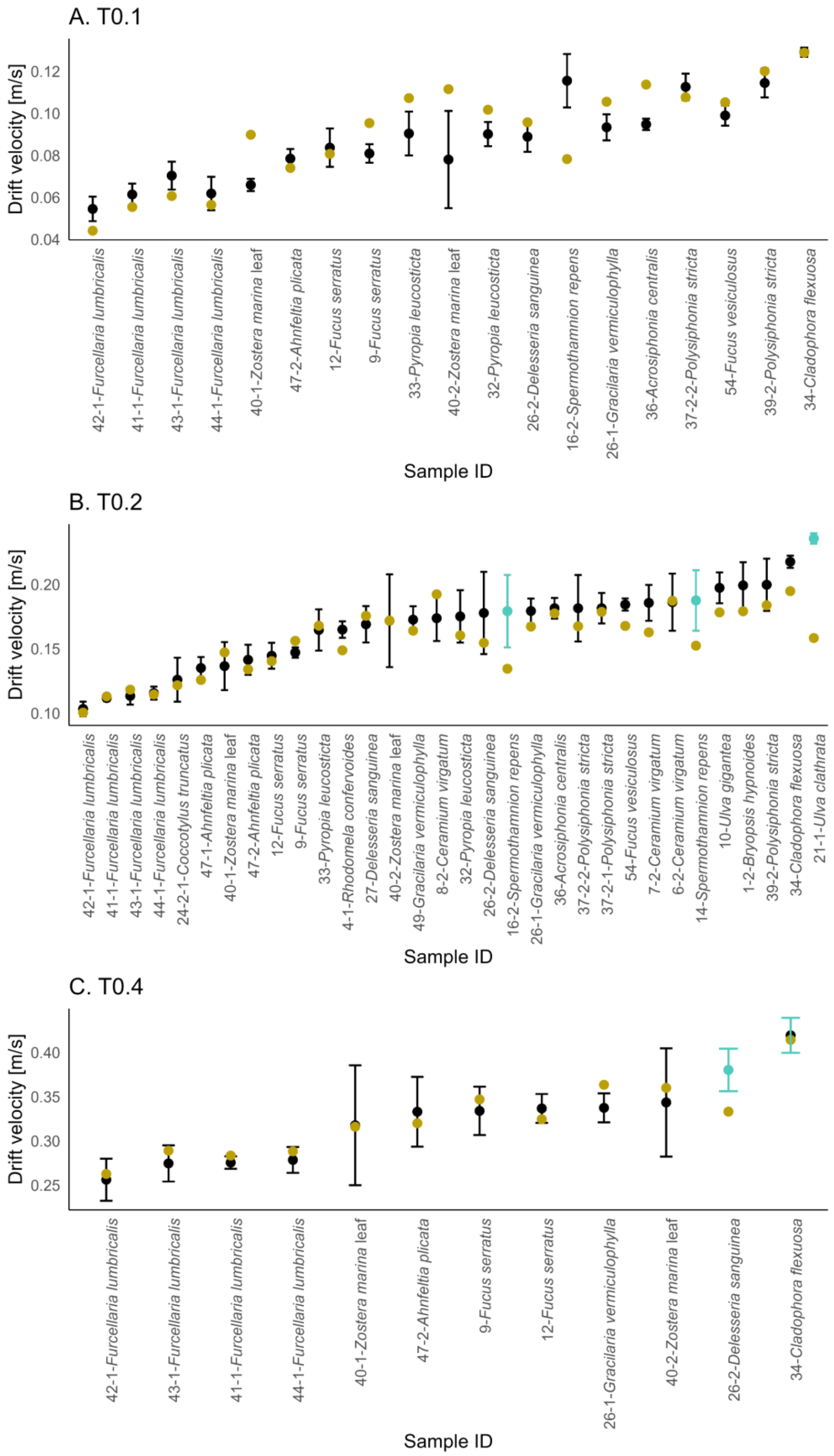


Supplementary Figure 10. Species specific model fit for negatively buoyant species using Equation 2. Error bars and points observed drift velocities, mean ± sd; black in contact with the bottom, blue drifting in the water column. Yellow dots modelled drift velocities.

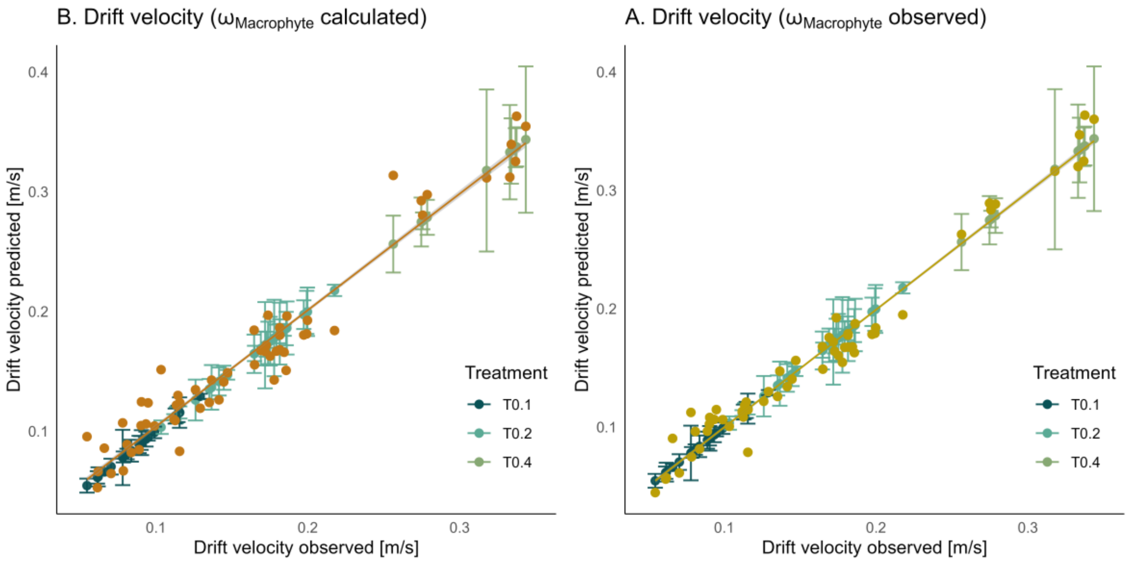


Supplementary Figure 11. Fit of the drift velocity model for negatively buoyant species, Equation 2. The observed drift velocities in blue and green (means ± sd) vs. the modelled drift velocities in yellow or orange (means). In Panel A., the observed macrophyte drag was used, whereas in Panel B., the calculated macrophyte drag was incorporated.
